# Functional characterisation of single nucleotide variants of the psychiatric risk gene *cacna1c* in the zebrafish

**DOI:** 10.1101/2021.09.30.462600

**Authors:** Nancy Saana Banono, Kinga Gawel, Tuomo Mäki-Marttunen, Wietske van der Ent, Wirginia Kukula-Koch, Marianne Fyhn, Gaute T. Einevoll, Ole A. Andreassen, Camila V. Esguerra

**Author notes:** Equally contributing (Kinga Gawel, Tuomo Mäki-Marttunen).

## Abstract

Several genome-wide association studies have associated *CACNA1C* variants with psychiatric disorders. The molecular mechanisms involved are poorly understood. Taking advantage of the zebrafish larva as a model, we investigated how two different mutations in *cacna1c – sa10930* (nonsense mutation) and *sa15296* (splice site mutation), affect neuronal function. We characterized changes in *cacna1c* mRNA, neurotransmitter levels and behaviour, as well as whole-brain activity using single electrode local field potential recordings. Both point mutations resulted in a significant reduction in *cacna1c* mRNA, as well as social behaviour and prepulse inhibition deficits. Whereas *sa15296* mutants displayed abnormal locomotor and open-field behaviour, we observed normal behaviour in the *sa10930* mutants. Brain recordings from both mutants had lower spectral power while *sa15296* displayed significant seizure-like activity. Finally, *sa10930* homozygotes showed increased dopamine and serotonin levels, decreased gamma-aminobutyric acid (GABA) levels, and unchanged glutamate levels while homozygous *sa15296* larvae showed increased levels of serotonin and glutamate, and unaffected levels of GABA and dopamine. Our work provides new insights into the functional role of *CACNA1C* in behavioural, electrophysiological and biochemical traits linked to psychiatric disorders. We show a functional role for the non-coding mutation (*sa15296*) in the *cacna1c in vivo* animal model. Consistent with existing hypotheses, our data suggest that disruption of gene expression, neurotransmission, and cortical excitability are involved in *CACNA1C*-related mechanisms of psychiatric disorders.

## Introduction

Neuropsychiatric disorders contribute to about 14% of the global burden of diseases [1]. Although the aetiology of these disorders is complex, involving the interplay of both genetic and environmental factors [2], their heritability is high. For schizophrenia (SCZ), bipolar disorder (BD) and autism spectrum disorders (ASD), heritability is estimated at up to 80% [3]. Many putative psychiatric susceptibility genes have been identified over the last decade *via* genome-wide association studies (GWAS) [4, 5]. One gene implicated by GWAS in mental disorders is *CACNA1C*, which encodes the alpha-1 subunit, Ca_V_1.2, of the L-type voltage-dependent calcium channel (LTCC) [6]. Although GWAS have implicated *CACNA1C* in mental disorders, the specific underlying molecular pathogenesis mechanisms are still unknown. We, therefore, investigated in this study, how mutations in *CACNA1C* affect psychiatry-related phenotypes *in vivo* using the zebrafish model.

*CACNA1C* is expressed in the brain, heart, smooth muscle, and endocrine cells [6, 7]. Neuronal Ca_V_1.2 channels are involved in gene expression regulation, neurotransmitter release, and integration of dendritic information in the brain [6–8]. In particular, neuronal Ca_V_1.2 are involved in several processes relevant to psychiatric disorders such as learning, memory, and brain development [7, 9].

Variations in *CACNA1C* have been associated with BD, SCZ and ASD, as well as other psychiatric disorders including attention deficit hyperactivity disorder (ADHD), and major depressive disorder [4, 5, 7]. How single nucleotide polymorphisms (SNPs) relate to molecular mechanisms associated with these disorders remains unclear. Interestingly, while some of the variants are within gene coding regions, most of the variants identified thus far are within non-coding regions [7, 10, 11]. The SNP rs1006737 confers a significant risk for BD, SCZ and ASD [12–15]. rs1006737 lies within intron 3 (a non-coding region) of the *CACNA1C* gene and is thought to be associated with altered gene expression [11, 16, 17]. The SNP rs4765905 is reported to affect interactions with the *CACNA1C* promoter region, thus, leading to an alteration in its expression [11]. Furthermore, two nonsense mutations in *CACNA1C* predicted to result in loss-of-function (LOF), have been identified through whole-exome sequencing in a SCZ population [10]. In addition, missense mutations in exons 8 or 8a of *CACNA1C* cause a rare subtype of ASD called Timothy syndrome^1^ *via* a gain-of-function (GOF) mechanism [9, 18]. Thus, there is a large interest in characterising the consequences of *CACNA1C* aberrations, as recently illustrated [19].

Although SCZ, BP and ASD are three distinct disorders, several studies have suggested that they have shared genetic risks, common pathologies and symptoms [20, 21]. The association of *CACNA1C* SNPs with multiple disorders suggests shared genetic convergence across disorders [5] and thus highlights the need to investigate the functional roles of *CACNA1C* using relevant disease endophenotypes rather than focusing on rigid disease classification.

Most of the functional roles of Ca_V_1.2 in psychiatric disorders have been obtained from genetic rodent models [9, 19] and to a lesser extent using zebrafish genetic models [22, 23]. Except for a recent aforementioned study [23], the role of *cacna1c* within the context of normal brain function, behaviour, and disease in zebrafish, has been underexplored. Furthermore, although most of the SNPs identified in humans are located in non-coding regions [11], there is no genetic animal model reported with mutations in non-coding regions of *Cacna1c*. The goal of the present study was to investigate how different mutations in *cacna1c* affect brain function in zebrafish larvae. Two mutant zebrafish lines generated *via* N-ethyl-N-nitrosourea (ENU) mutagenesis were obtained from the zebrafish international resource centre. The first line, *sa10930*, results in a premature stop codon in exon 6, whereas the second line, *sa15296*, carries an essential splice site mutation in intron 35 [24]. To understand the relationship between the gene variants and central nervous system (CNS) function, we examined the morphological, molecular, behavioural, electrophysiological, and biochemical characteristics of mutant larvae *versus* their wild type (WT) counterparts. Together, these comprehensive experiments provide new knowledge on how variants of *cacna1c* may contribute to psychiatric disease.

## Materials and methods

### Zebrafish mutant lines and husbandry

Two mutant lines *cacna1c ^sa10930^* (hereafter *sa10930* for homozygotes and *sa10930/WT* for heterozygotes) and *cacna1c ^sa15296^* (hereafter *sa15296* for homozygotes and *sa15296/WT* for heterozygotes) that were generated by ENU mutagenesis from the Zebrafish Mutation Project (Sanger, UK) were obtained as fertilised embryos from the Zebrafish International Resource Center (Eugene, Oregon, USA) and raised to adulthood.

Adult zebrafish stocks were maintained under standard conditions [25] in an approved fish facility with a 14 hours’ light and 10 hours’ dark cycle. Fertilised eggs from the natural spawning of adult fish lines were collected, transferred to petri dishes (n = 60), and raised in an incubator at 28°C in E3 medium. All the experiments were approved by the Norwegian Food Safety Authority experimental animal administration’s supervisory and application system (FOTS-18/106800-1; ID 15469 and 23935).

### Genotyping

The allelic-specific primers: *sa15296* forward 5’-AAGACTGTGGCAGTCACTTTG-3’, *sa15296* reverse 5’-ACTGTACGGAGGGGGTAAAA-3’, *sa10930* forward 5’ - ATGGTTCCCCTCCTTCAC-3’ and *sa10930* reverse 5’ - AGTTCAAGGGAGAAGCAAAAG-3 ‘ were used under cycling conditions shown in (**tables 1 & 2**). The restriction enzymes, MseI and HphI were used to digest the PCR products obtained from the *sa10930* and *sa15296* lines respectively.

**Table 1:** PCR cycling conditions for *sa10930*

| Step | Temperature °C | Time | Number of cycles |
| --- | --- | --- | --- |
| Initial denaturation | 95 | 30 sec | 1 |
| Denaturation | 95 | 30 sec | 35 |
| Annealing temperature | 57 | 30 sec |  |
| Extension | 72 | 1 min |  |
| Final extension | 72 | 1 min | 1 |

**Table 2:** PCR cycling conditions for *sa15296*

| Step | Temperature °C | Time | Number of cycles |
| --- | --- | --- | --- |
| Initial denaturation | 95 | 30 sec | 1 |
| Denaturation | 95 | 30 sec | 35 |
| Annealing temperature | 57.5 | 30 sec |  |
| Extension | 72 | 1 min |  |
| Final extension | 72 | 1 min | 1 |

### Morphological analysis

Live anaesthetised larvae were mounted using 2% methylCellulose. Images of larvae were taken with the Leica M205 FA microscope at the same resolution for direct comparison.

### Mannitol exposure

Larvae were exposed to 250 mM mannitol (#M4125, Sigma) as described in [26] from 3 days post-fertilization (dpf) and replenished daily until the last day larvae were used for experiments. The plates housing larvae were changed after every two days to avoid solute crystals that form due to evaporation of the solution. 3 dpf was chosen as the treatment start day because oedema was visible from this day onwards.

### Whole-mount *in-situ* hybridization (WISH)

WISH was used to investigate the spatial gene expression patterns of *cacna1c* using digoxigenin labelled riboprobes as earlier described [27]. Briefly, 5 and 7 dpf larvae were fixed in 4% paraformaldehyde in 1 × PBS. The mMESSAGE mMACHINE™ SP6 Transcription Kit, mMESSAGE mMACHINE™ T7 Transcription Kit (Thermo Scientific Fisher), and DIG RNA labeling Mix (Roche) were used to make the Digoxigenin (DIG) UTP-labeled RNA riboprobes. The final concentration of cacna1c probes used was 200 ng in 200 μl at a staining duration of 35 minutes. Images of larvae stained with the sense and antisense probes were taken using the iDS camera (in optimal colours mode) mounted on a Zeiss Stemi 508 stereomicroscope. Below are the primer sequences for the sense and antisense probes:

cacna1c_sense+T7_Fwd: TAATACGACTCACTATAGGG TCAGAAGACA GCACTACCAT
cacna1c_sense_Rev: TGGGCTAATACACAGGACGT
cacna1c_antisense_Fwd: TCAGAAGACA GCACTACCAT
cacna1c_antisense+T7_Rev: TAATACGACTCACTATAGGG GGGCTAATACACAGGACGT

### Quantitative reverse transcription (rt-qPCR)

6 dpf homozygous *sa15296, sa10930*, and their respective WT siblings were collected (25 larvae/sample, n = 3/group) for the extraction of RNA. Total RNA from larvae flash frozen in liquid nitrogen was isolated using Invitrogen PureLink RNA Mini Kit (12183018A, Thermofischer). The quality of RNA was checked using a NanoDrop 1000 Spectrophotometer (Thermofischer) and the agarose bleach gel method as described previously [28]. cDNA was synthesized from 0.5 μg RNA templates using the SuperScript First-Strand Synthesis System for rt-PCR (Invitrogen) as described by the manufacturer. The cDNA was diluted by a factor of 60. The rt-qPCR was performed in triplicates with 9 μl of the diluted cDNA and 2× PowerUp Sybr Green Master Mix (Applied Biosystems) in a 20 μl final volume. The 96-well plate was used under the following cycling conditions: 95 °C for 10 min, 94 °C for 15 s, 60 °C for 60 s for 40 cycles followed by dissociation analysis using 65 to 95 °C at 0.5 °C increments. *Glyceraldehyde-3-phosphate dehydrogenase* (*gapdh*) was used as the reference gene [see primer sequences below].

*gapdh_Fwd:* 5’ GTGGAGTCTACTGGTGTCTTC 3’
*gapdh_Rev:* 5’ GTGCAGGAGGCATTGCTTACA 3’
*cacna1c (transcript variants 201 and 202)*_Fwd: 5’ GGATAACGCCAGGATCTCAA 3’
*cacna1c (transcript variants 201 and 202)*_Rev: 5’ AGCAGAACAGCCTGAGGAAA 3’
*cacna1c (transcript variant 202)*_Fwd: 5’ GACCCCTGGAATGTTTTTGA 3’
*cacna1c (transcript variant 202)*_Rev: 5’ CATTGGTCTTACCACAGAGGAG 3’

### Western blotting

Homozygous *sa15296, sa10930*, and their respective WT siblings at 7 dpf were collected (20 larvae/sample, n = 3/group) in a master mix containing RIPA buffer and halt protease inhibitor cocktail (Thermo Fischer) for the extraction of immune-blot lysate. Samples were sonicated (30 s run, 30 s pause for ten times at 4 °C), centrifuged for 10 minutes at 4 °C, the supernatant collected and an equal volume of 2X laemmli buffer added and stored in - 20 °C until ready to use. There was no boiling of lysate. Protein was separated using Mini-PROTEAN TGX Precast Gel (#4569033EDU, Bio-rad) and transferred on to a nitrocellulose membrane at 30 V for 16 hours. Blocking was done in 5 % milk powder for 1 hour at room temperature. Incubation in the primary antibodies i.e Ca_V_1.2 antibody (monoclonal, 1: 1000, #MA5-27717, Thermofischer), anti-Ca_V_1.3 antibody - C-terminal (3:100, #ab191038, Abcam) and monoclonal mouse anti-β actin antibody (1: 1000, #MA1-744, Thermofischer) at 4 °C overnight. Incubation in the secondary antibodies i.e. polyclonal goat anti-rabbit conjugated horseradish peroxidase (1: 20000, #31460, Thermofischer) and polyclonal goat anti-mouse conjugated horseradish peroxidase (1: 20000, #G-21040, Thermofischer) for 2 hours at room temperature. SuperSignal West Pico PLUS Chemiluminescent Substrate (Thermofischer) was used for protein visualization. The total Cacna1c protein level was obtained after it was normalized relative to the β-actin protein level.

Additionally, the anti-Ca_V_1.2 antibody (polyclonal, 1:200, #ab58552, Abcam) was also tested. Both anti-Ca_V_1.2 antibodies (#MA5-27717 from Thermofischer and #ab58552 from Abcam) recognize largely the same areas within the c-terminal of cacna1c. The c-terminal of cacna1c is conserved between human and zebrafish.

### Behavioural analyses

The ZebraBox (ViewPoint, France) was used for video recording of larvae to determine locomotor, thigmotaxis and shoaling behaviour whereas the ZebraBox Revo (ViewPoint, France) with PPI add-ons was used for the prepulse inhibition (PPI) test. ZebraLab (ViewPoint, France) and EthoVision (Noldus, Netherlands) software were used to analyse larval locomotor activity. On the day of the experiments, the fish were transported from the incubator, allowed to acclimate to the conditions of the recording room for a minimum of one hour. All recording sessions were conducted between 10:00 to 18:00 hours during the day and under a temperature-controlled setting (27±1°C).

#### Locomotor activity

For measuring locomotor activity, larvae (6 dpf) were acclimated 15 min in the ZebraBox (testing chamber) in a 48 well plate under the same illumination conditions (light or dark) as the tracking conditions followed by a 10 minutes recording session in a 1 min time bin. The parameters measured were the average total distance travelled in millimetres (mm) and the duration spent in inactivity in seconds (s).

#### Light-dark transition test

For the light-dark transition test, larvae (6 dpf) were acclimated in the test chamber for 15 minutes followed by 10 min each of tracking in (1) darkness (0 % light) 2) 100% light, and (3) darkness. Larval locomotor activity was measured as the total distance moved (mm) over either a 10 min period in time bins of 1 min or a complete recording period.

#### Thigmotaxis

Thigmotaxis was measured using a 24 well plate (diameter = 16.2 mm) as the open field. Each well of the plate was divided into an outer and inner zone (diameter of inner zone = 8 mm, inner zone distance in relation to the outer zone = 4 mm) and thigmotaxis was calculated as the total distance moved (TDM) in the outer zone as described by [29, 30]. The preference of each fish to remain at the periphery or explore the centre of the arena was monitored for 20 minutes of spontaneous behaviour in either light or dark conditions. Larvae aged 6 dpf were acclimated 15 min in the test chamber before the start of recording.

#### Shoaling

The shoaling assay was performed using a round dish of diameter 60 mm, height 15 mm, and volume 30 ml (82.1194.500, Sarstedt). Larvae (7 dpf) were gently placed in the center of the testing arena (5 larvae/arena, n = 10-14/group) and acclimated for 15 min followed by a 20 minute recording session at 30s-time bin to obtain a time-series data using the ZebraLab software. The following social dynamics: nearest neighbor distance (NND) and inter-individual distance (IID) of the shoal were measured as previously described [31, 32].

#### Acoustic startle response (ASR) and PPI

The acoustic startle response and PPI of 6 dpf larvae were evaluated using the ZebraBox Revo (ViewPoint, France) and EthoVision (Noldus, Netherlands) as we previously showed [33]. Single larvae were placed in individual wells of a custom-made plexiglass plate (33-wells in a 96 format). Larvae were acclimated in a 100 Lx illuminated ZebraBox 5 min before the onset of the experiment. A 660 Hz startle stimulus of duration 100 ms and 440 Hz prepulse stimulus of duration 5 ms were used. The inter-stimulus interval (ISI) for all PPI experiments was 100 ms.

### Local field potential (LFP) analysis

The LFP recordings of 7 dpf larvae were performed as previously described [34]. A glass electrode was filled with artificial cerebrospinal fluid and inserted into the optic tectum of individual larvae aged 7 dpf, immobilised using a thin layer of 2% low melting point agarose. Whole-brain activity was measured for 20 minutes using a MultiClamp 700B amplifier and Digidata 1550 digitizer (Axon Instruments, USA). The Clampfit version 10.6.2 software (Molecular Devices Corporation, USA) was used for processing the LFP recordings. The data were analysed manually by two independent trained observers, blind to the genotype of the larvae.

For spectral analysis, the LFP time series were divided into time windows of 10 seconds with a 50% overlap, and the spectral power was determined for each segment using Welch’s method as implemented in the python package scipy.stats. The spectrograms were visually inspected, and the fish that displayed a prominent peak at 2.5 Hz, which we interpreted as a heart-beat component, were excluded from the analysis: this resulted in discarding 4 recordings from WT larvae and 2 recordings from each of the mutant larvae. Moreover, within each sample, the time windows where the total spectral power was more than twice the median spectral power across the time windows were discarded, along with two neighbouring time windows before and after the high-power window. This was done to remove transient artefacts. The remaining time windows were pooled across the fish of the same genotype. This resulted in 1838 spectral samples for the WT larvae, 1644 samples for the *sa10930* larvae, and 1394 samples for the *sa15296* larvae. Each spectral sample was then normalized by the total spectral power within the range 0.5-200 Hz after filtering out the power-grid component at 50 Hz and its duplicates (100, 150 Hz) with a margin of ±0.5 Hz. To compare the mutant fish spectra with the WT spectra, Wilcoxon rank-sum test (U-test) was used with a threshold p-value of 0.01, which was Bonferroni-corrected by the number of frequency windows (980) to 1.02 × 1 0-5.

For determining the slope of the spectral power decay, we transformed the data (both frequency and power) into a logarithmic scale and performed a linear regression within the frequency range 40 - 200 Hz, filtering out the power-grid components as above. The distributions of the slopes were compared with each other using the U-test with a threshold p-value of 0.01.

### Sample preparation and neurotransmitter analysis

7 dpf larvae were collected (100 larvae/sample, n = 4-10 per group) in a tube, excess fish medium removed, then snap-frozen in liquid nitrogen. Cold acetonitrile (stored at −20°C) and milli-Q water were added at a volume of 200 μl each. Sonication at 4 °C and centrifugation at 24000 rpm for 5 min. The supernatant was collected in clean Eppendorf tubes for further analysis. HPLC-MS grade acetonitrile, water, and formic acid were purchased from Merck Millipore (Darmstadt, Germany). Dopamine hydrochloride (#PHR1090), serotonin hydrochloride (#H9523), γ-aminobutyric acid (#03835), and glutamic acid (#49449) purchased from Sigma Aldrich were used as standards.

Quantitative analysis of gamma-aminobutyric acid (GABA), glutamate, dopamine and serotonin were performed using the HPLC-ESI-Q-TOF-MS/MS platform as previously described [35]. The HPLC-ESI-QTOF-MS/MS platform produced by Agilent Technologies (Santa Clara, CA, USA) was used for the development of chromatographic methods and the studies on the composition and quantification of the selected neuromodulators based on the accurate mass measurements, the fragmentation patterns, and in consideration of the data from open databases (Metlin). The instrument was composed of a 1200 Series HPLC chromatograph with a degasser, binary pump, an autosampler, a column oven, a photodiode array detector DAD and an ESI-QTOF-MS/MS mass spectrometer (G6530B). The platform was calibrated in the positive and negative ionization modes before the study. Both chromatographic and spectrometric conditions were initially optimized for each solution of the reference compound to obtain in terms of gradient composition and mass spectrometer settings to receive clearly separated peaks and an elevated response from the MS spectrometer (gas temperatures, capillary voltage, collision energy settings). Accucore HILIC HPLC column (150 x 2.1 mm, 2.6 μm particle size) by Thermo Fischer Scientific (Waltham, MA, USA) was used in the study.

The following 70-minute long gradient program with 0.2 mL/min flow rate of 0.1% formic acid in water (solvent A) and 0.1% formic acid in acetonitrile (solvent B) was applied: 0 min 98% of B, 5 min 80% of B, 40 min 75% of B, 45 min 30% of B, 48 min 98% of B. The method was recorded for 60 min. A 10 μL injection, 25°C oven temperature, 10 min long post run were used. The mass spectrometer was operated under the following mild conditions to provide higher sensitivity to the acquisition of small molecules like the tested neuromodulators: gas and sheath gas temperatures: 275 and 300 °C, respectively, the gas flows: 12 L/min, nebulizer voltage: 35 psig, capillary voltage: 3000 V, fragmentor voltage: 80 V, collision energy: 5 and 10 V, m/z detection range: 50-1200 Da. In the method, after the collection of one MS/MS spectrum given signals were excluded for the following 0.2 min from further fragmentation to obtain the MS/MS spectra of further less intensive peaks. Agilent Mass Hunter Workstation (v. B.08.00) was used for the acquisition of spectra and the analysis of recorded data.

### Statistical analysis

Version 8.4.1 of the GraphPad Prism software (San Diego, CA, USA) was used for statistical analyses and generation of graphs except for power spectral analysis of the LFP data. Unpaired Students *t*-test or its equivalent non-parametric test, Mann Whitney was used in comparing two groups. Where more than two groups were compared, one-way analysis of variance (ANOVA) followed by Tukey’s *post-hoc* test was performed. We used two-way ANOVA followed by Tukey’s *post-hoc* test to analyse the light-dark transition test data and to determine the effects of stimulus type on the PPI assay. Mann-Whitney U-test and Kruskal-Wallis test followed by Dunn’s *post-hoc* tests were used to calculate the statistics for thigmotaxis and the light-dark tests respectively. Statistical significance was established when p < 0.05 except for spectral power analysis where p < 0.01.

## Results

### Molecular and morphological effects induced by *cacna1c* mutations

The *sa10930* mutant line carries a specific thymidine (T) to adenosine (A) point mutation in exon 6 (NM_131900.1: c.876T>A: p.Tyr292Stop), which results in a premature stop codon (TAA) at amino acid 292 (**Fig. 1a**). *In silico* analysis predicts exon 6 to form part of the first transmembrane domain that regulates voltage sensitivity of the Ca_V_1.2 channel (**supplementary Fig. S1**). Thus, premature termination may result in channel loss of function. On the other hand, the *sa15296* mutant line carries a thymidine (T) to cytosine (C) point mutation in the donor splice site between intron 35-36 (NC_007115.7: g.155870T >C), resulting in an essential splice variant (**Fig 1a**). We predict that the *sa15296* mutation may result in exon skipping and/or nonsense-mediated mRNA decay. Either of the aforementioned situations could alter the sensitivity of the channel – whether it leads to overall hypo/hyperactive channel activity is yet to be determined.

**Figure 1.**
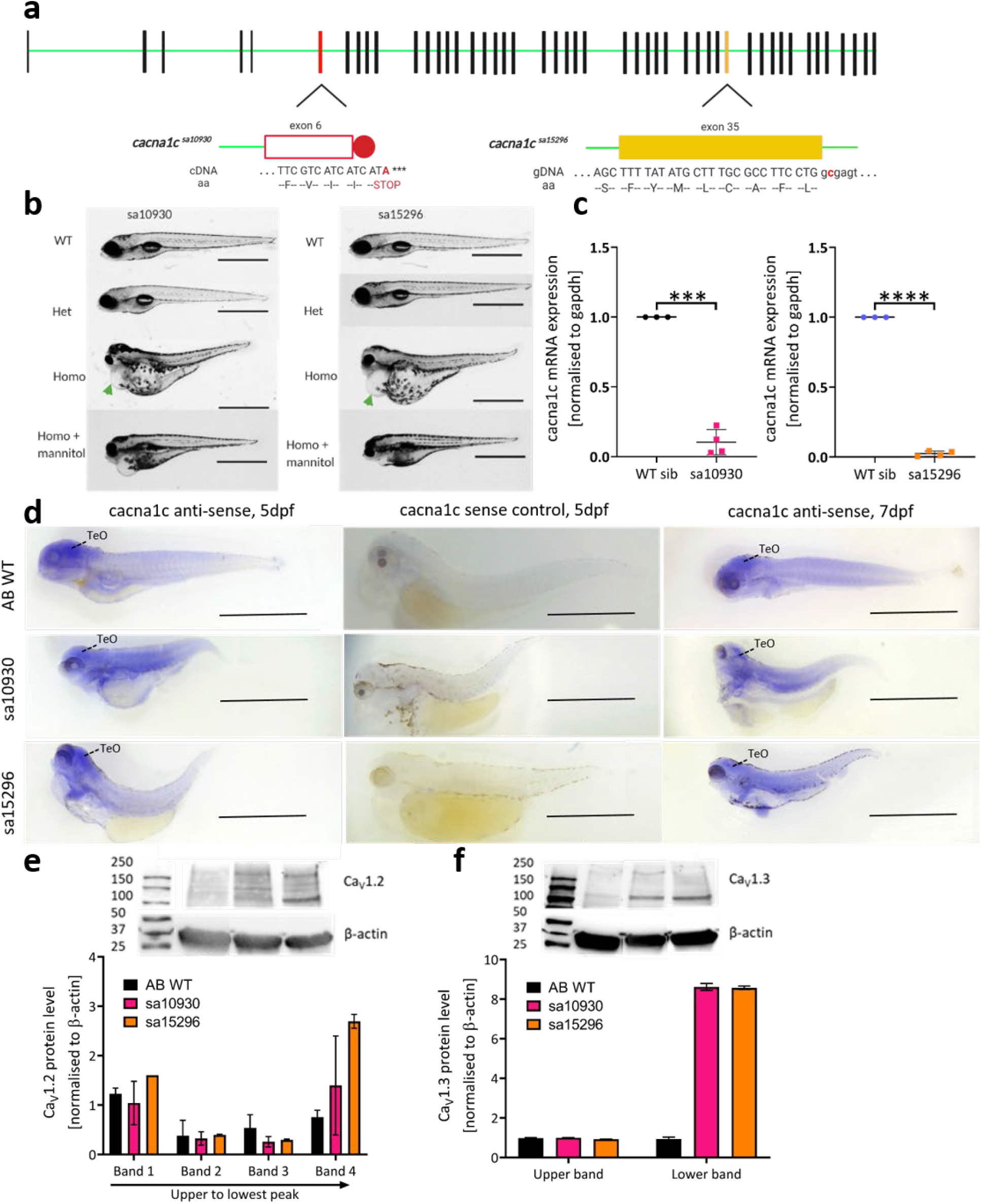
Zebrafish *cacna1c* mutant lines and their molecular and morphologic effects on larvae. a) Schematic representation of zebrafish *cacna1c* based on the sequence from ensembl genome (ENSDART00000028856.5 (GRCz11)), not drawn to scale. There is a total of 47 exons depicted as solid vertical bars and introns shown as green lines. The colours red and orange are used to highlight the region of *sa10930* and *sa15296* mutation respectively. Nucleotides in upper case are within the exon while those in lower case are within the intron. Round red ball represents protein truncation. b) Morphology of mutants at 5 dpf. Heterozygous *sa10930* and *sa15296* mutant larvae are morphologically indistinguishable from their WT siblings. Homozygous *sa10930* and *sa15296* mutant larvae have pericardial oedema (green arrow), show hyperpigmentation and craniofacial abnormalities compared to siblings, scale bar = 750 μm. WT – wild-type, Het – heterozygote, Homo – homozygote. c) Representative whole mount *in situ* hybridization with antisense and sense probes of *cacna1c*. Dpf, days post-fertilization; TeO, optic tectum d) rt-qPCR analysis revealed reduced expression of *cacna1c* in homozygous (d) *sa10930* [t (19.82) = 3, (p < 0.001); welch corrected] and (e) *sa15296* [t (105.7) = 3, (p < 0.0001); welch corrected] mutants at 7 dpf. Data was analysed using unpaired Student *t*-test and represented as mean ± S.D. For analysis, n = 3-4 samples/group. e) Representative western blot image and analysis of Cacna1c protein. Bands 1-4 correspond to noticeable bands in a descending order of protein size (biggest to smallest). f) Representative western blot image and analysis of Cacna1da protein. Bands 1-2 correspond to noticeable bands in a descending order of protein size (biggest to smallest).

To determine the genotype of our experimental animals, restriction enzyme digestion of the PCR products was performed. The *sa10930* mutation introduces an MseI restriction site into the *cacna1c* gene. Digestion of the 161 base pair (bp) long PCR product spanning the mutation with MseI results in two smaller products (109 and 52 bp) in fish carrying the mutation - i.e., two bands for homozygous, three bands for heterozygous, and one band for WT after digestion (**supplementary Fig. S2a**). On the contrary, the *sa15296* mutation did not introduce a novel restriction site. However, the WT sequence already harboured an HphI restriction site. Digestion of the 181 bp long PCR product spanning the area of the mutation with HphI results in two smaller products (112 and 69 bp) in non-mutant fish i.e. one band for homozygous, three bands for heterozygous, and two bands for WT after digestion (**supplementary Fig. S2b**).

When heterozygous mutants of either *sa15296* or *sa10930* were in-crossed, the genotypes of the resulting offspring were within the expected Mendelian proportion of 25% WT, 50% heterozygous mutant, and 25% homozygous mutant (data not shown). Heterozygous mutants were morphologically indistinguishable from their WT siblings, were viable, fertile, and survived to adulthood. After 48 hpf, both *sa15296* and *sa10930* homozygous mutants showed several developmental abnormalities such as oedema of the pericardial region, yolk sac and gut that worsened with age, as well as craniofacial abnormalities and curved body axis (**Fig. 1b**). We observed no deficits in touch response of both *sa10930* and *sa15296* mutants in either the heterozygous or homozygous state. Since the mutations were larval lethal, homozygous mutants of the two lines did not survive past 10 dpf.

Our interest in performing a repertoire of behavioural tests across genotypes motivated us to identify ways to reduce oedema severity in homozygous mutants, as this impaired locomotor ability. We achieved this by increasing the osmolarity of the surrounding environment (**Fig. 1b**). Maintaining larvae in 250 mM mannitol from 3 dpf significantly decreased the severity but did not prevent the occurrence of oedema or prolong the life span of the mutants. Importantly, mannitol did not affect larval locomotor activity, as this could potentially serve as a confounding factor in interpreting behavioural responses (**supplementary. Fig. S3**).

At the gene transcription level, rt-qPCR analysis at 7 dpf revealed a significant reduction in *cacna1c* mRNA levels by at least 90 % of homozygous *sa10930* [t (19.82) = 3.000, p < 0.001; Welch-corrected] (**Fig. 1c**) and *sa15296* [t (105.7) = 3.000, p < 0.0001; Welch-corrected] when compared to WT using *gapdh* as the reference gene (**Fig. 1c**).

We examined the spatiotemporal expression of *cacna1c* transcript(s) in homozygous *sa10930* and *sa15296* as well as WT larvae at 5 and 7 dpf, using WISH (**Fig. 1d**). In WT larvae, *cacna1c* was expressed in the heart, brain and to a lesser extent in the musculature, which is consistent with previous reports [36]. Overall, there was a low expression of *cacna1c* in the brain of homozygous *sa10930* and *sa15296* mutant larvae at 5 dpf with reduced expression sustained up to 7 dpf (the latest developmental stage tested in our experimental setup). There was no visible expression of *cacna1c* in larvae stained with the sense probes.

Next, we examined whether the decreased expression of *cacna1c* mRNA in the two homozygous mutant lines was reflected in a change in protein levels. We performed immunostaining using 7 dpf zebrafish whole-lysate with anti-Ca_V_1.2 antibody (**Fig. 1e**). Interestingly, there was a comparable to a slight increase in the signal of anti-Ca_V_1.2 antibody in both the null mutant (*sa10930*) and the splice variant mutant (*sa15296*) relative to WT. The result was surprising in that the anti-Ca_V_1.2 antibody recognizes the c-terminus of Cacna1c. Since the *sa10930* mutant harbours a truncated channel lacking the c-terminus, we expected no anti-Ca_V_1.2 signal whereas a significantly low amount of anti-Ca_V_1.2 signal was expected in the *sa15296* mutant at least based on the WISH and rt-qPCR data, which showed that the *cacna1c* transcript levels in both mutants were reduced. We evaluated the similarity of the antibody sequence with Cacna1d^2^, another LTCC to confirm any cross-reactivity or otherwise. Indeed, we observed significant homology of the anti-Ca_V_1.2 antibody sequence to that of Cacna1da upon protein blast analysis (**supplementary Fig. S4a**). Next, we performed immunostaining using anti-Ca_V_1.3 antibody shown to have significantly less sequence homology with Cacna1c (**supplementary. Fig S4b**). The results revealed a significant increase in anti-Ca_V_1.3 signal in both homozygous *sa10930* and *sa15296* mutants relative AB WT (**Fig. 1f**).

### Effects of *cacna1c* mutations on larval behaviour

To assess the impact of *cacna1c* mutations on larval locomotor behaviour, larvae (6 dpf) were tracked for 10 minutes under dark conditions. Kruskal-Wallis H test revealed a statistically significant difference between larvae [*sa10930*: average total distance (H = 52.56, p < 0.0001) (**Fig. 2a**) and average duration of inactivity (H = 6.227, p < 0.05)] (**supplementary Fig. S5a**) and [*sa15296*: average total distance (H = 58.34, p < 0.0001) (**Fig. 2b**) and the average duration of inactivity (H = 9.088, p < 0.05)] (**supplementary Fig. S5b**). Dunn’s *post-hoc* test revealed that *sa10930* homozygous larvae displayed a statistically significant decrease in locomotor activity (p < 0.0001) and increased inactivity duration (p < 0.0001) in relation to WT and heterozygous *sa10930* siblings. Heterozygous *sa10930* mutants had comparable locomotor activity and inactivity duration with WT siblings (p > 0.05). On the other hand, reduced locomotor activity was observed in both heterozygous (p < 0.05) and homozygous *sa15296* (p < 0.0001) when compared to their WT siblings. Whilst no difference in the duration spent in inactivity between heterozygous *sa15296* and WT siblings (p > 0.05) was seen, homozygous *sa15296* spent more time being inactive (p < 0.0001) when compared to both heterozygous mutants and WT siblings.

**Figure 2.**
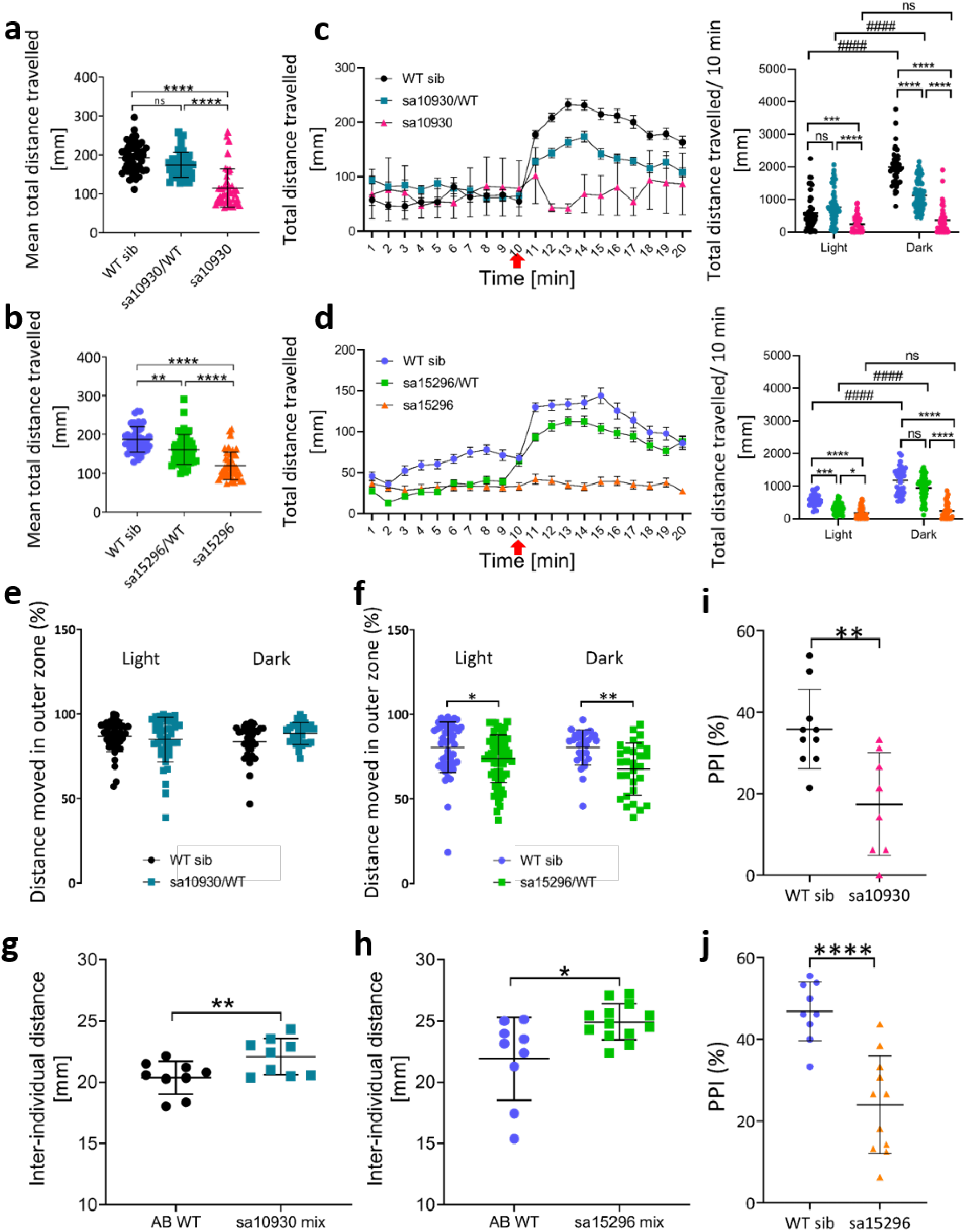
Behavioural effects of *cacna1c* mutations on zebrafish larvae. a-b) Locomotor activity of *cacna1c* mutants versus their WT siblings shown as the average total distance travelled over 10 min. (a) *sa10930* and (b) *sa15296*. Data was analysed using Kruskal-Wallis one-way ANOVA test followed by Dunn’s multiple comparison test and represented as mean ± S.D. For the analysis, n = 42-48, **p < 0.01, ****p < 0.0001. c-d) Response of larvae to abrupt changes in illumination in the light-dark transition test. (c) *sa10930* and (d) *sa15296*. Data represented as mean ± S.E.M for line graph and Mean ± S.D. for column graph. Analysis was done using the Kruskal Wallis test followed by Tukey’s multiple comparison test *p < 0.05, ***p < 0.001, ****p < 0.0001, #### p < 0.0001, ^^^^ p < 0.0001. The black arrow indicates lights turned on and the grey shadow from 10 min onwards indicate lights turned off. e-f) Zone preference of 6 dpf larvae under dark and light conditions in the thigmotaxis test. Thigmotaxis was measured as the % total distance moved in the periphery of the arena. (e) There was no difference in the zone preference of the *sa10930* mutants. (f) *sa15296* mutants had a significant preference for the centre of the arena than their WT siblings under both light and dark conditions. Data was analysed using the Mann-Whitney U test and represented as mean ± S.D. For the analysis, n = 27-33, *p < 0.05, **p < 0.01. g-h) Increased inter-individual distance exhibited by a heterogeneous population of heterozygous mutants and their wild-type siblings versus homogeneous wild-type population. (g) *sa10930* mix and (h) *sa15296* mix. Data was analysed using unpaired Student *t*-test and represented as mean ± S.D. For the analysis, n = 9-13 (5 larvae/shoal), *p < 0.05, **p < 0.01. i-j) Impaired prepulse inhibition of the acoustic startle response of homozygous mutant larvae at 6 dpf in the sensorimotor test. (i) *sa10930* and (j) *sa15296*. Data was analysed using unpaired Student *t*-test and represented as mean ± S.D. For the analysis, n = 8-16, ** p < 0.01, **** p < 0.0001.

We evaluated if mutants displayed anxiety-like behaviour by subjecting them to the light-dark test which utilises abrupt changes in illumination from light to dark to elicit startle and/or stress responses, as measured by an exaggerated locomotor activity [37-39] and the open field test where we measured thigmotaxis. The normal response for larvae in a light-dark and dark-light switch is hyperlocomotion and freezing respectively [38, 39]. Whereas heterozygous mutants of both lines showed a normal response in the dark-light and light-dark switch, homozygous mutants of both lines remained unresponsive to either switch. However, the extent of the response of heterozygous *sa10930* in the light-dark switch was lesser than WT [(p < 0.0001), **Fig. 2c**] relative to WT. For the *sa15296* group, heterozygotes exhibited profound freezing in the light [(p < 0.001), **Fig. 2d**] when compared to WT. We further analysed the behaviour of mutants in the open field test, a measurement of novelty-associated anxiety. Heterozygous *sa10930* mutants showed no significant difference in zone preference under both dark (U = 319, p > 0.05) and light (U = 889, p > 0.05) conditions, although there was a tendency of mutants to stay in the outer zone when compared with their WT counterparts (**Fig. 2e**). On the contrary, heterozygous *sa15296* mutants spent significantly more time in the inner zone in both dark (U = 211, p < 0.001) and light (U = 1291, p < 0.01) conditions than their WT siblings (**Fig. 2f**) indicating boldness or risk-taking behaviour [40].

Furthermore, because persons with disorders such as SCZ and ASD present social deficits [21], we tested if the *cacna1c* mutants also displayed social deficits by subjecting them to a shoaling^3^ test. We measured two shoaling characteristics – nearest neighbour distance (NND)^4^ and interindividual distance (IID)^5^. The *sa10930* mix group exhibited impaired social behaviour as evidenced by increases in NND [t (20) = 2.184, (p < 0.05); (**supplementary Fig. S5c**)] and IID [t (20) = 2.860, p < 0.01; (**Fig. 2g**)] when compared with their WT siblings. Although the *sa15296* mix group showed no difference in NND [t (16) = 0.5639, p > 0.05; (**supplementary Fig. S5d**)], we observed a significant increase in IID [t (16) = 2.547, p < 0.05] (**Fig. 2h**) in relation to their WT siblings, which is also suggestive of impaired social behaviour.

Finally, we examined whether our mutants harboured any sensorimotor deficits by exposing them to acoustic startle stimuli and measuring the extent of PPI. The zebrafish, like many animals, has an innate startle response that is attenuated when preceded by a weak, non-startling stimulus [41]. Similar to humans and rodents, the pharmacological agents apomorphine, haloperidol and ketamine modulate the zebrafish PPI thus supporting its translational value [33, 41]. Presentation of a prepulse did not attenuate the startle response of both homozygous *sa10930* (**Fig. 2i**) and *sa15296* (**Fig. 2j**) mutants as evidenced by their decreased % PPI when compared to their WT counterparts.

### Effects of *cacna1c* mutations on the whole-brain activity of zebrafish larvae

We measured the spectrograms of homozygous mutants of the two *cacna1c* lines and WT fish and pooled the time-windowed spectra across the fish of the same genotype (see Methods). We compared the power spectra of the mutant fish with those of WT in the delta (1 – 4 Hz), theta (4 – 8 Hz), alpha (8 – 13 Hz), and beta (13 – 30 Hz) frequency ranges. We determined the LFP slope by performing linear regression in the log-log scale within the gamma (30 –100 Hz) frequency band. The spectra of all fish showed large intrinsic variability (**Fig. 3a-c**). When we averaged the time-windowed data for each fish instead of the pooling procedure, the differences were non-significant both for the individual spectral power components (U-test, p > 0.05, Bonferroni corrected) and for the spectral slopes (U-test, p > 0.05). However, when pooled across time windows, homozygous fish from both *cacna1c* mutant genotypes showed a significantly lower spectral power in the low delta-frequency (0.5-2 Hz) range than WT (**Fig. 3d**, U-test, p<0.01, Bonferroni-corrected). In other frequency ranges, the mutant fish only showed sporadic differences in the beta and gamma bands (**Fig. 3d**). The spectral slopes, determined for the frequency range 50 – 200, were significantly flatter in the mutants than in the WT fish (**Fig. 3e**), suggesting increased excitability in the neural circuits of the mutant fish [42]. Taken together, these results show that the *cacna1c* mutants, despite a large intrinsic variability, display systematic LFP-power deficits in the low delta range and the spectral slopes.

**Figure 3.**
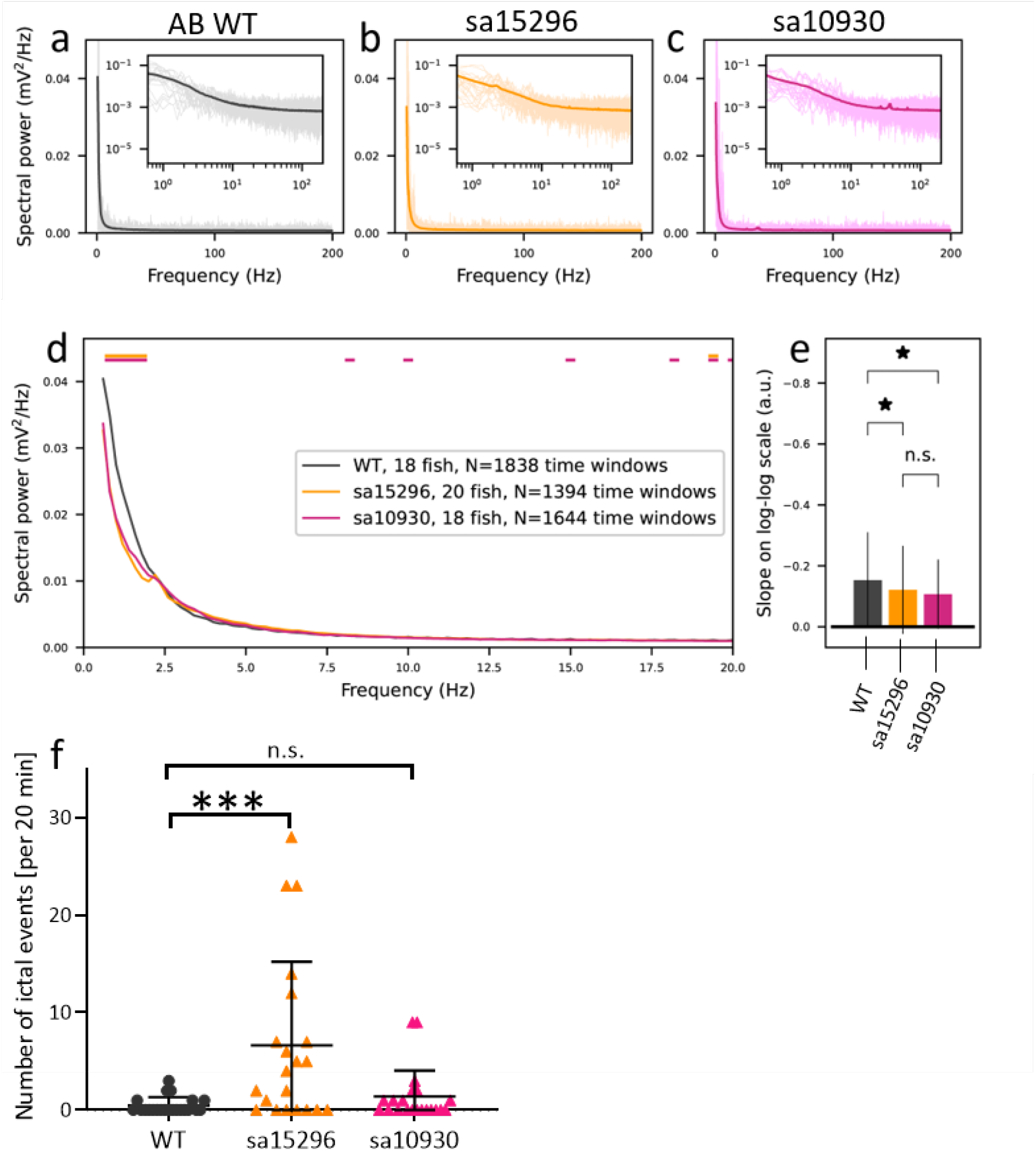
Delta-range EEG power and the spectral slope are reduced in *cacna1c* mutants compared to WT larvae. a-c) EEG spectra of 10-second EEG time series of a) WT, b) *sa15296* mutant and c) *sa10930* mutant larvae. d) Dim curves show 20 randomly picked samples, and the thick lines show the pooled averages. Insets show the same data in the log-log scale. The bars above the curves show the frequency components where the power of the mutant larvae (*sa15296* for the upper bars, *sa10930* for the lower bars) differed significantly (p<0.01) from the corresponding frequency component of the WT larvae. e) The mean pooled spectra of panels a-c overlaid and zoomed in on 0-20 Hz. The bars show the mean spectral slope of each fish genotype and the lines show the SD. Both mutant fish types had a significantly flatter slope (p<0.01; the numbers of samples were the same as in panel (D)). f) Number of ictal discharges observed in mutants larvae during the 20 minutes LFP recording session. Data analysed using one-way ANOVA followed by the Dunnet’s *post-hoc* test and represented as mean ± S.D.. For the analysis, n = 21-22, *** p < 0.001. WT – wild-type, Homo – homozygote.

Further examination of the LFP recordings revealed seizure-like discharges in mutant larvae. One-way ANOVA analysis indicated statistically significant differences concerning the number of seizure-like discharges [F (2, 61) = 10.92, p < 0.001]. The *sa15296* mutants discharged more seizure-like activity p < 0.001 when compared to WT larvae whereas there was no statistically significant difference between the *sa10930* mutants and WT larvae (**Fig. 3f**). Sample traces of seizure-like activity of WT and mutants are shown in (**supplementary Fig. S6**).

### Effects of *cacna1c* mutations on the neurotransmitter levels

Imbalances in neurotransmitter levels have long been hypothesised to be associated with the pathoaetiology of SCZ and BD [43, 44]. We used HPLC analysis to assess if the neurotransmitter levels of glutamate, GABA, dopamine, and serotonin in the mutants were altered relative to their WT siblings at 7 dpf (**supplementary Table T1 & supplementary Fig. 7**).

Increased levels of both dopamine (p < 0.05) and serotonin (p < 0.01) and decreased GABA levels (p < 0.0001) were observed, while the levels of glutamate were unaffected (p > 0.05) in the homozygous *sa10930* mutants (**Fig. 4a**). However, in the homozygous *sa15296* mutants we observed an increased amount of both glutamate and serotonin at p < 0.05, while the levels of both GABA and dopamine remained unaffected at p > 0.05 (**Fig. 4b**). Further analysis of the glutamate/GABA ratio, a measure of excitation – inhibition, revealed that both mutant lines had significantly increased glutamate/GABA [(WT *vs sa10930/sa10930;* p < 0.01) (**Fig. 4c**) and (WT *vs sa15296/sa15296;* p < 0.05)] (**Fig. 4d**).

**Figure 4.**
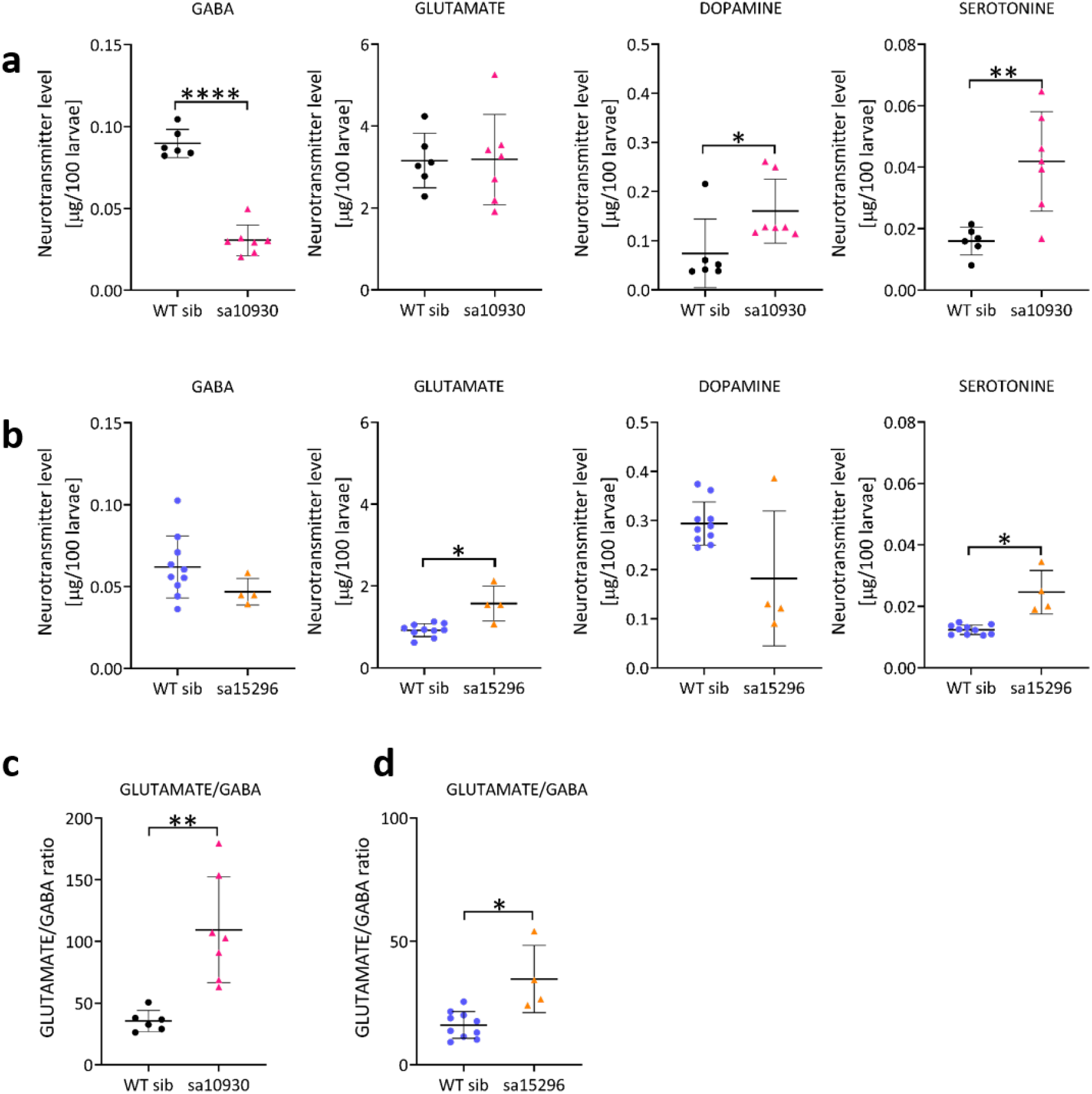
Mutations in *cacna1c* lead to imbalances in the neurotransmitter levels of glutamate, GABA, dopamine and serotonin. a) *sa10390*, b) *sa15296*, c) Glutamate/GABA ratio of WT vs *sa10930* and d) Glutamate/GABA ratio of WT *vs sa15296*. Data was analysed using unpaired Student *t*-test and represented as mean ± S.D. For the analysis, n = 4 – 10, *p < 0.05, **p < 0.01, and ****p < 0.0001.

### Effects of *cacna1c* mutations on downstream molecular targets

Calcium influx through Ca_V_1.2 calcium channels regulates downstream genetic transcription pathways such as brain-derived neurotrophic factor (BNDF) and c-Fos [45, 46]. We investigated the influence of low levels *cacna1c* on *bndf* and *c-fos* to find possible molecular pathways that may be responsible for the observed phenotypes so far described. We found a statistically significant reduction in the mRNA levels of *bdnf* (p < 0.05) but not *c-fos* (p > 0.05) in homozygous *sa10930* mutants relative to WT siblings (**Fig. 5a**). However, in the homozygous *sa15296* mutants, the mRNA levels of both *bdnf* (p < 0.05) and *c-fos* (p < 0.05) were statistically significant when compared to their WT siblings (**Fig. 5b**).

**Figure 5.**
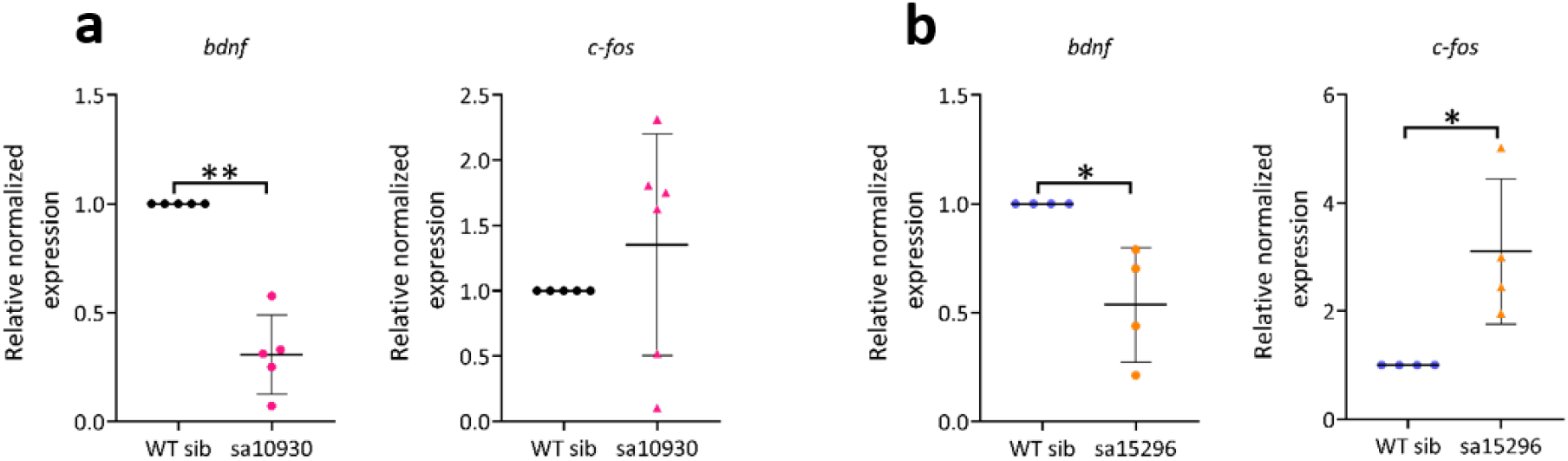
Mutations in *cacna1c* lead to changes in selected downstream targets. a) Changes in *bdnf* and *c-fos* expression in homozygous *sa10930* mutants. b) Changes in bdnf and c-fos expression in homozygous *sa15296* mutants. Data was analysed using unpaired Student *t*-test and represented as mean ± S.D. For the analysis, n = 4 – 10, *p < 0.05, **p < 0.01, and ****p < 0.0001.

## Discussion

Here, we investigated the molecular mechanisms affecting brain function of the psychiatric risk gene *CACNA1C* in genetically modified zebrafish larvae. We characterized behavioural, electrophysiological and biochemical traits linked to psychiatric disorders in two mutant lines harbouring point mutations in distinct regions (i.e., coding (*sa10930*) and non-coding (*sa15296*) regions) of the zebrafish *cacna1c* gene to gain further information on the functional role of CACNA1C gene variants in brain disorders.

The *sa10930* mutation is a nonsense mutation that results in a premature stop codon in exon 6 and reduced gene expression in homozygotes. We found no evidence of exon skipping as a result of the *sa15296* mutation although it occurs in the essential splice site. Rather, we observed reduced expression of cacna1c mRNA. Similarly, previous studies identified a non-coding SNP that resulted in reduced *CACNA1C* expression [11, 16, 17]. The data thus suggests LOF in both mutants and indicates autoregulation of *cacna1c* expression. Furthermore, our data show that *cacna1c* LOF results in a GOF of *cacna1da*, through a compensatory upregulation of cacna1da in our fish models. Nevertheless, the morphological phenotype (pericardial oedema, craniofacial abnormalities) and larval lethality (survival < 10 dpf) of homozygous *sa10930* and *sa15296* mutants is similar to what has been reported for both *cacna1c* LOF zebrafish [23, 36, 47] and rodents [48, 49].

Behavioural analysis of the two mutant lines revealed both overlapping and distinct phenotypes. Locomotor defects (decreased swimming distance and increased duration of inactivity) were evident in homozygous *sa10930* and homozygous *sa15296*, similar to what was observed in earlier studies involving LOF *cacna1c* zebrafish larvae [23], likely as a direct result of the oedema present in these fish. Although heterozygous mutants did not show any morphological impairments, heterozygous *sa15296* exhibited decreased locomotor activity while their *sa10930* counterparts showed similar activity when compared to WT larvae.

With exception of homozygous *sa10930* and *sa15296* mutants, all other larvae were responsive to abrupt changes in illumination. In the dark to light switch test, the swimming distance of heterozygous *sa10930* was unaffected while that of heterozygous *sa15296* was significantly reduced when compared to their respective WT controls. We observed no changes in zone preference in heterozygous *sa10930* larvae under both dark and light conditions, whereas the opposite was the case for heterozygous *sa15296* larvae. A previous study showed that although zebrafish *cacna1c* mutants (LOF) have a normal response to light stimuli, they tend to stay at the periphery of an open field (thigmotaxis) as well as display an abnormal response to darkness i.e., absence of light [23]. The aforementioned phenotypes i.e., thigmotaxis and abnormal response to darkness are reminiscent of anxiety-like behaviours in zebrafish [29, 38]. Notably, several studies have also reported increased anxiety in mouse models of *Cacna1c* dysfunction [50-53]. We excluded homozygous mutants from thigmotaxis measurements because the quantification of thigmotaxis relies on adequate locomotor capabilities [29]. Taken together, these findings suggest an effect on behaviour that may have implications for mental traits and disorders.

Zebrafish are social animals that are attracted to conspecifics and tend to move in a group [31, 32]. Although the attraction of larvae to conspecifics starts as early as 6 dpf, shoaling has been shown to develop with age but is evidenced as early as 7 dpf, albeit less robust than at older stages [54]. In zebrafish, the shoaling assay is commonly used in validating models for ASD and to a lesser extent, models for SCZ and intellectual disability [55]. By analysing two shoaling parameters, deficits in NND and IID were found in *sa10930* mixed genotype larvae while for *sa15296* mixed genotype larvae, deficits were only observed in the IID. It is important to note that NND is independent of shoal size while IID is dependent on shoal size. Hence, some researchers consider IID reflective of shoal cohesion as opposed to NND [31, 56]. However, measuring both provide further information about the shoal dynamics. We did not use homozygous mutants in the shoaling test because of their high immobility, which could bias the results. Furthermore, it was impossible to segregate heterozygous mutants from their WT counterparts since they are morphologically indistinguishable. Thus, by taking advantage of the “heterogeneous shoaling paradigm” previously described [57], mixed populations of fish (i.e., a non-homogeneous genotype group made up of heterozygous mutants and WT siblings) were used for the shoaling assay.

If a weaker auditory stimulus (prepulse) precedes a startle stimulus within a period of 30-500 ms, the magnitude of response to the startle stimulus is suppressed in a dose-dependent manner with respect to the prepulse strength. This phenomenon, referred to as PPI, is a quantitative measure of sensorimotor gating and happens already at the first trial, hence, no learning is required. Sensorimotor gating is representative of the brain’s ability to differentiate between relevant and irrelevant sensory input to make a proper behavioural response [58, 59]. Reduced PPI is considered a useful behavioural endophenotype for SCZ in human and animal models [41, 60, 61]. In SCZ and other psychiatric disorders and their equivalent animal models, deficits in PPI have been reported [41, 62]. Thus, defective PPI is indicative of information processing dysfunction [58]. We observed reduced PPI responses in homozygous larvae carrying both *sa10930* and *sa15296* mutations. This observation is similar to what was earlier reported for *cacna1c* LOF zebrafish larvae [23] and for humans carrying mutations in other neuropsychiatric related genes [59]. The brain circuitry and neurotransmitter systems mediating PPI in mammals are conserved in the zebrafish [63, 64].

We observed a significant reduction in the expression of total *bdnf* in mutants *versus* WT using whole larval isolate. There is evidence of reduced *Bdnf* expression in conditional *Cacna1c* conditional knockout mice [65], and region-specific (↓ *Bdnf* in the prefrontal cortex, ↑ *Bdnf* in the dentate gyrus of the hippocampus) regulation of *Bdnf* in the *Cacna1c* heterozygous rats [66]. In humans, risk variants in *CACNA1C* were associated with decreased expression in *CACNA1C* and as well as region-specific BDNF regulation (↓ *BDNF* in the prefrontal cortex and substantia nigra, ↑ *BDNF* in the dentate gyrus of the hippocampus) [66]. Bdnf deficient mice showed enhanced acoustic startle response (ASR), reduced PPI of ASR and deficits in nesting behaviour (a social behaviour) [58].

Analysis of LFP spectrograms from mutant and WT larval brains was performed to test whether *cacna1c* mutant larvae display electrophysiological phenotypes that correspond to mental disorder-associated LFP-based phenotypes. Delta-range spectral power and spectral slope, have been shown to be altered in SCZ and ADHD, respectively [67, 68]. Both of the *cacna1c* mutants showed a significantly lower spectral power in the low delta-frequency (0.5-2 Hz) range than WT. In other frequency ranges, mutant larvae only showed sporadic differences in the beta and gamma band. The spectral slopes, determined for the frequency range 40 - 200 Hz, were significantly flatter in the mutant than in WT, suggesting increased excitability in the neural circuits of the mutants [42]. Hyper-excitability of neural circuitry underlies a number of brain disorders, particularly those associated with seizure-like activity [42, 69]. Indeed, seizure-like discharges were observed in *sa15296* but not *sa10930* mutants. The discharges observed in *sa15296* mutants generally have a lower amplitude and a shorter duration compared to the EEG patterns of the pentylenetetrazole-induced seizure model or the genetic Dravet model [34, 69]. Recent studies have revealed pathogenic variants in *CACNA1C* associated with epilepsy, some of which involve splice site mutations [70]. The clinical spectrum of *CACNA1C* variants is highly variable, likely due to a variety of reasons, such as the nature of the mutation - i.e., its location within the protein and structural domain of Ca_V_1.2 and the varying transcript expression in different tissues [70].

The increased neural excitability of mutants suggested by the less negative slopes from the spectral analysis was further supported by 1) the increased glutamate/GABA ratio for both *sa10930* and *sa15296* mutants relative to their WT siblings from neurotransmitter analysis and 2) increased levels *c-fos* mRNA. Imbalances in neurotransmitters, in particular dopamine, have long been hypothesized to be associated with the aetiology of mental disorders [43, 44]. Disruption in GABA-, glutamate-, and dopaminergic neurotransmission is thought to contribute to perturbed cortical oscillations [71]. A recent paper identified risk genes associated with a wide range of neurotransmitter systems - glutamatergic, GABAergic, dopaminergic, serotonergic, cholinergic and opioid in SCZ [44]. We observed enhanced dopamine and serotonin, decreased GABA, and unaffected levels of glutamate in homozygous *sa10930* larvae. In the homozygous *sa15296* larvae, enhanced serotonin and glutamate, unaffected levels of GABA, and dopamine were observed. Calcium channel activity modulates overall levels of neurotransmitters through the activation of G-protein coupled receptors which control the sensitivity of ionotropic neurotransmitter receptors and ion channels [44]. Therefore, mutations in calcium channel genes can alter neuronal excitability and firing patterns possibly through alterations in neurotransmission. Notably, however, neurotransmitter levels were analysed from whole-larvae in our experimental setup and therefore does not provide information regarding tissue-specific changes.

The observed reduced expression of *bdnf* in *sa10930 and sa15296* larvae suggests that decreased *cacna1c* alters downstream signalling pathways such as *bdnf* that may be mediating the observed phenotypes herein reported.

Our multi-trait analyses provide interesting insights into the functional role of *CACNA1C* in psychiatric disorders using LOF mutants. Our earlier computational modelling studies predicted that LOF *CACNA1C* variants can lead to 1) decreased “single-cell” PPI in pyramidal cells [72, 73], 2) decreased cardiac activity [74], and 3) increased delta oscillations (and overall activity) in networks of pyramidal neurons [72]. The present results lend support to the first prediction but are inconsistent with the third prediction. The reason for the incongruency with the predictions on delta-oscillation power in a mammalian cortex may be the large density of SK currents in the mammalian L5 pyramidal cells, which are likely to cause gain-of-function mutations of high-voltage-activated Ca2+ channels to counter-intuitively decrease the neuronal excitability in these cells, and *vice versa* [74]. It is not known whether the neuronal types giving rise to the LFP signal in the delta frequency range in zebrafish larvae express SK currents. Previous studies of *Cacna1c* mutations in rodents [75] and clinical samples [76] indicated that LOF mutations were associated with altered sleep quality. Moreover, LOF mutations of CACNA1C decreased high-frequency LFP oscillations during wake and REM sleep in patients [75], where a (non-significant) decrease of low-frequency delta-power can also be observed. Further research is needed to increase our understanding of the effects of CACNA1C mutations on delta oscillations and sleep. As for the second prediction, our finding of decreased mobility of homozygous *cacna1c* mutant larvae (as a consequence of severe pericardial oedema), may be related to reported cardiac deficits [77], but other causes such as rest behaviour, anxiety-induced freezing behaviour or neuromotor defects are also possible [40].

Even though we confirmed that the mutations harboured by the *sa10930* and *sa15296* resulted in over 90% decrease (LOF) of *cacna1c* at the mRNA level, we could not replicate the *cacna1c* LOF finding at the protein level due to the non-specificity of the anti-Ca_V_1.2 antibodies used. The substantial amount of anti-Ca_V_1.2 signal detected especially in the *sa10930* mutant samples could not be Cacna1c because the mutation causes a truncated channel lacking the c-terminus region, which the epitope recognizes. However, considering the high homology of the anti-Ca_V_1.2 epitope to the c-terminus of Cacna1d, it is likely that the observed protein signal is Cacna1d. Upon further analysis, we found the protein level of Cacna1d to be significantly increased when immune-blot lysates were probed with antibodies specific for Cacna1d. These results, altogether, suggest a compensatory upregulation of Cacna1d following *cacna1c* LOF. Although *CACNA1D* is the most likely LTCC to compensate for a LOF of *CACNA1C* [78, 79], earlier studies conducted in Cacna1c mice with 1) global happloinsufficiency [51], 2) conditional knockout in neurons [80] and 3) conditional deletion in the hippocampus and neocortex [81] were reported to have unaffected Cacna1d protein levels. The disparity could stem from the fact that the presence of one healthy Cacna1c allele is enough to support essential functions hence no need for any compensatory expression by “sister” proteins while the timing and/or regions of the conditional deletion could be outside critical windows and/or areas that drive the need for any compensatory expression. Notably, CACNA1D GOF mutations have been linked to neurodevelopmental disorders such as ASD and epilepsy [82-84]. Whether the reported phenotypes in this study and for the other reported studies are solely a result of Cacna1c LOF, of Cacna1d GOF, or a combination of both is presently unclear and warrants further investigation.

Our work provides additional insights into the putative functional role of CACNA1C using a number of behavioural, electrophysiological and biochemical traits linked to psychiatric disorders. More specifically, we show for the first time, a functional role of a non-coding *cacna1c* mutation (*sa15296*) in an *in vivo* animal model. Taken together, our data provide additional evidence that *cacna1c* LOF 1) disrupts ion channel activity-dependent autoregulation of gene expression, 2) dysregulates neurotransmission and 3) increases cortical excitability. Such perturbations during early brain development result in altered behaviours reminiscent of those described for persons with psychiatric disorders, including an altered response to dark and light, anxiety-like behaviour, reduced social interaction and an impaired PPI response. However, whether all observed phenotypes are a result of *cacna1c* LOF and/or *cacna1da* GOF needs to be investigated further, both in fish and in rodent CACNA1C and CACNA1D models.

## Author contribution

NSB, CVE, KG, TM, OAA, GTE and MF conceptualised the experiments. NSB carried out all larval morphological assessment, behavioural phenotyping, rt-qPCR, western blotting, HPLC sample preparation and corresponding data analysis. KG and WE performed LFP experiments and seizure analysis. TM performed spectral analysis of LFP data. WK-K carried out HPLC measurement and analysis. NSB, KG and TM prepared the figures. All authors contributed to the preparation of the manuscript. NSB and CVE revised and edited the final version. All authors read and approved the manuscript.

## Acknowledgements

We thank Ana Tavara and Alejandro Pastor Remiro for fish care.

Open Access funding provided by University of Oslo and Oslo University Hospital. This work was funded by the Research Council of Norway [RCN# 248828; (ISP, BIOTEK2021/ DigiBrain), for NSB and CVE] and the Centre for Molecular Medicine Norway start-up funds (for CVE) and RCN grant 223273 (for OAA). KG received funding from the European Union’s Horizon 2020 research and innovation programme (Marie Skłodowska-Curie, No. 798703-GEMZ-H2020-MSCA-IF-2017).

## List of main abbreviations

ADHD: attention deficit hyperactivity disorder
ASD: autism spectrum disorders
ASR: acoustic startle response
BD: bipolar disorder
BDNF: brain-derived neurotrophic factor
Dpf: days postfertilization
ENU: N-ethyl-N-nitrosourea
GABA: gamma-aminobutyric acid
GOF: gain-of-function
GWAS: genome-wide association studies
Hpf: hours post-fertilization
IID: interindividual distance
LFP: local field potential
LOF: loss-of-function
LTCC: L-type voltagedependent calcium channel
NND: nearest neighbour distance
PPI: prepulse inhibition
SNP: single nucleotide polymorphism
WISH: whole-mount *in-situ* hybridization
WT: wild type

## Footnotes

1 Timothy syndrome (TS) is an autosomal dominant disease characterized by autism, impaired cognitive functions, craniofacial abnormalities and cardiac defects [18].

2 The CACNA1D gene is duplicated in zebrafish into cacna1da and cacna1db. Cacna1da is 77 % while cacna1da is 33 % homologous to the human orthologue [ensembl.org].

3 Shoaling is the tendency of fish to remain in close proximity to conspecifics [31].

4 NND measures the distance between the nearest neighbours within a shoal [31].

5 IID measures the average distance among all members of a shoal [31].

## References

1. Prince M, Patel V, Saxena S, Maj M, Maselko J, Phillips MR, et al. No health without mental health. The Lancet. 2007;370:859–877.

2. Patel KR, Cherian J, Gohil K, Atkinson D. Schizophrenia: overview and treatment options. P T. 2014;39:638–645.

3. Winchester CL, Pratt JA, Morris BJ. Risk genes for schizophrenia: translational opportunities for drug discovery. Pharmacol Ther. 2014;143:34–50.

4. Ripke S, Neale BM, Corvin A, Walters JT, Farh K-H, Holmans PA, et al. Biological Insights From 108 Schizophrenia-Associated Genetic Loci. Nature. 2014;511:421–427.

5. Cross-Disorder Group of the Psychiatric Genomics Consortium. Identification of risk loci with shared effects on five major psychiatric disorders: a genome-wide analysis. The Lancet. 2013;381:1371–1379.

6. Heyes S, Pratt WS, Rees E, Dahimene S, Ferron L, Owen MJ, et al. Genetic disruption of voltage-gated calcium channels in psychiatric and neurological disorders. Prog Neurobiol. 2015;134:36–54.

7. Kabir ZD, Martínez-Rivera A, Rajadhyaksha AM. From Gene to Behavior: L-Type Calcium Channel Mechanisms Underlying Neuropsychiatric Symptoms. Neurotherapeutics. 2017;14:588–613.

8. Greer PL, Greenberg ME. From Synapse to Nucleus: Calcium-Dependent Gene Transcription in the Control of Synapse Development and Function. Neuron. 2008;59:846–860.

9. Moon AL, Haan N, Wilkinson LS, Thomas KL, Hall J. CACNA1C: Association With Psychiatric Disorders, Behavior, and Neurogenesis. Schizophr Bull. 2018;44:958–965.

10. Purcell SM, Moran JL, Fromer M, Ruderfer D, Solovieff N, Roussos P, et al. A polygenic burden of rare disruptive mutations in schizophrenia. Nature. 2014;506:185–190.

11. Roussos P, Mitchell AC, Voloudakis G, Fullard JF, Pothula VM, Tsang J, et al. A Role for Noncoding Variation in Schizophrenia. Cell Reports. 2014;9:1417–1429.

12. Ferreira MAR, O’Donovan MC, Meng YA, Jones IR, Ruderfer DM, Jones L, et al. Collaborative genome-wide association analysis supports a role for ANK3 and CACNA1C in bipolar disorder. Nat Genet. 2008;40:1056–1058.

13. Li J, Zhao L, You Y, Lu T, Jia M, Yu H, et al. Schizophrenia Related Variants in CACNA1C also Confer Risk of Autism. PLoS One. 2015;10.

14. Nyegaard M, Demontis D, Foldager L, Hedemand A, Flint TJ, Sørensen KM, et al. CACNA1C (rs1006737) is associated with schizophrenia. Mol Psychiatry. 2010;15:119–121.

15. Sklar P, Smoller JW, Fan J, Ferreira MAR, Perlis RH, Chambert K, et al. Whole-genome association study of bipolar disorder. Mol Psychiatry. 2008;13:558–569.

16. Gershon ES, Grennan K, Busnello J, Badner JA, Ovsiew F, Memon S, et al. A rare mutation of CACNA1C in a patient with bipolar disorder, and decreased gene expression associated with a bipolar-associated common SNP of CACNA1C in brain. Mol Psychiatry. 2014;19:890–894.

17. Yoshimizu T, Pan JQ, Mungenast AE, Madison JM, Su S, Ketterman J, et al. Functional implications of a psychiatric risk variant within CACNA1C in induced human neurons. Mol Psychiatry. 2015;20:162–169.

18. Napolitano C, Antzelevitch C. Phenotypical manifestations of mutations in the genes encoding subunits of the cardiac voltage-dependent L-type calcium channel. Circ Res. 2011;108:607–618.

19. Tigaret CM, Lin T-CE, Morrell ER, Sykes L, Moon AL, O’Donovan MC, et al. Neurotrophin receptor activation rescues cognitive and synaptic abnormalities caused by hemizygosity of the psychiatric risk gene Cacna1c. Mol Psychiatry. 2021. 17 February 2021. https://doi.org/10.1038/s41380-020-01001-0.

20. Guan J, Cai JJ, Ji G, Sham PC. Commonality in dysregulated expression of gene sets in cortical brains of individuals with autism, schizophrenia, and bipolar disorder. Transl Psychiatry. 2019;9:152.

21. Vorstman JAS, Burbach JPH. Autism and Schizophrenia: Genetic and Phenotypic Relationships. In: Patel VB, Preedy VR, Martin CR, editors. Comprehensive Guide to Autism, New York, NY: Springer New York; 2014. p. 1645–1662.

22. Banono NS, Gawel K, De Witte L, Esguerra CV. Zebrafish Larvae Carrying a Splice Variant Mutation in cacna1d: A New Model for Schizophrenia-Like Behaviours? Mol Neurobiol. 2021;58:877–894.

23. Thyme SB, Pieper LM, Li EH, Pandey S, Wang Y, Morris NS, et al. Phenotypic Landscape of Schizophrenia-Associated Genes Defines Candidates and Their Shared Functions. Cell. 2019;177:478–491.e20.

24. Kettleborough RNW, Busch-Nentwich EM, Harvey SA, Dooley CM, de Bruijn E, van Eeden F, et al. A systematic genome-wide analysis of zebrafish protein-coding gene function. Nature. 2013;496:494–497.

25. Aleström P, D’Angelo L, Midtlyng PJ, Schorderet DF, Schulte-Merker S, Sohm F, et al. Zebrafish: Housing and husbandry recommendations: Laboratory Animals. 2019. 11 September 2019. https://doi.org/10.1177/0023677219869037.

26. Hill AJ, Bello SM, Prasch AL, Peterson RE, Heideman W. Water permeability and TCDD-induced edema in zebrafish early-life stages. Toxicol Sci. 2004;78:78–87.

27. Thisse C, Thisse B. High-resolution in situ hybridization to whole-mount zebrafish embryos. Nat Protoc. 2008;3:59–69.

28. Aranda PS, LaJoie DM, Jorcyk CL. Bleach gel: A simple agarose gel for analyzing RNA quality. ELECTROPHORESIS. 2012;33:366–369.

29. Schnörr S, Steenbergen P, Richardson M, Champagne D. Measuring thigmotaxis in larval zebrafish - ScienceDirect. 2012. https://www.sciencedirect.com/science/article/pii/S0166432811008758?via%3Dihub. Accessed 5 December 2019.

30. Liu X, Zhang Y, Lin J, Xia Q, Guo N, Li Q. Social Preference Deficits in Juvenile Zebrafish Induced by Early Chronic Exposure to Sodium Valproate | Behavioral Neuroscience. 2016. https://www.frontiersin.org/articles/10.3389/fnbeh.2016.00201/full. Accessed 5 December 2019.

31. Miller N, Gerlai R. Quantification of shoaling behaviour in zebrafish (Danio rerio). Behavioural Brain Research. 2007;184:157–166.

32. Miller N, Gerlai R. From Schooling to Shoaling: Patterns of Collective Motion in Zebrafish (Danio rerio). PLoS ONE. 2012;7:e48865.

33. Banono NS, Esguerra CV. Pharmacological Validation of the Prepulse Inhibition of Startle Response in Larval Zebrafish using a Commercial Automated System and Software. JoVE. 2020:61423.

34. Afrikanova T, Serruys A-SK, Buenafe OEM, Clinckers R, Smolders I, de Witte PAM, et al. Validation of the zebrafish pentylenetetrazol seizure model: locomotor versus electrographic responses to antiepileptic drugs. PLoS ONE. 2013;8:e54166.

35. Gawel K, Kukula-Koch W, Banono NS, Nieoczym D, Targowska-Duda KM, Czernicka L, et al. 6-Gingerol, a Major Constituent of Zingiber officinale Rhizoma, Exerts Anticonvulsant Activity in the Pentylenetetrazole-Induced Seizure Model in Larval Zebrafish. IJMS. 2021;22:7745.

36. Rottbauer W, Baker K, Wo ZG, Mohideen M-APK, Cantiello HF, Fishman MC. Growth and Function of the Embryonic Heart Depend upon the Cardiac-Specific L-Type Calcium Channel α1 Subunit. Developmental Cell. 2001;1:265–275.

37. Basnet RM, Zizioli D, Taweedet S, Finazzi D, Memo M. Zebrafish Larvae as a Behavioral Model in Neuropharmacology. Biomedicines. 2019;7:23.

38. Kedra M, Banasiak K, Kisielewska K, Wolinska-Niziol L, Jaworski J, Zmorzynska J. TrkB hyperactivity contributes to brain dysconnectivity, epileptogenesis, and anxiety in zebrafish model of Tuberous Sclerosis Complex. Proc Natl Acad Sci USA. 2020;117:2170–2179.

39. Li F, Lin J, Liu X, Li W, Ding Y, Zhang Y, et al. Characterization of the locomotor activities of zebrafish larvae under the influence of various neuroactive drugs. Ann Transl Med. 2018;6:173.

40. Kalueff AV, Gebhardt M, Stewart AM, Cachat JM, Brimmer M, Chawla JS, et al. Towards a Comprehensive Catalog of Zebrafish Behavior 1.0 and Beyond. Zebrafish. 2013;10:70–86.

41. Burgess HA, Granato M. Sensorimotor gating in larval zebrafish. J Neurosci. 2007;27:4984–4994.

42. Gao R, Peterson EJ, Voytek B. Inferring synaptic excitation/inhibition balance from field potentials. NeuroImage. 2017;158:70–78.

43. Sigitova E, Fišar Z, Hroudová J, Cikánková T, Raboch J. Biological hypotheses and biomarkers of bipolar disorder: Hypotheses of bipolar disorder. Psychiatry Clin Neurosci. 2017;71:77–103.

44. Devor A, Andreassen O, Wang Y, Mäki-Marttunen T, Smeland O, Fan C-C, et al. Genetic evidence for role of integration of fast and slow neurotransmission in schizophrenia. Mol Psychiatry. 2017;22:792–801.

45. Ghosh A, Carnahan J, Greenberg M. Requirement for BDNF in activity-dependent survival of cortical neurons. Science. 1994;263:1618–1623.

46. West AE, Griffith EC, Greenberg ME. Regulation of transcription factors by neuronal activity. Nat Rev Neurosci. 2002;3:921–931.

47. Ramachandran KV, Hennessey JA, Barnett AS, Yin X, Stadt HA, Foster E, et al. Calcium influx through L-type Ca_V_1.2 Ca^2+^ channels regulates mandibular development. J Clin Invest. 2013;123:1638–1646.

48. Kisko TM, Braun MD, Michels S, Witt SH, Rietschel M, Culmsee C, et al. *Cacna1c* haploinsufficiency leads to pro-social 50-kHz ultrasonic communication deficits in rats. Dis Model Mech. 2018;11:dmm034116.

49. Seisenberger C, Specht V, Welling A, Platzer J, Pfeifer A, Kühbandner S, et al. Functional Embryonic Cardiomyocytes after Disruption of the L-type α _1C_ (*Ca _v_ 1.2*) Calcium Channel Gene in the Mouse. J Biol Chem. 2000;275:39193–39199.

50. Bader PL, Faizi M, Kim LH, Owen SF, Tadross MR, Alfa RW, et al. Mouse model of Timothy syndrome recapitulates triad of autistic traits. Proceedings of the National Academy of Sciences. 2011;108:15432–15437.

51. Dao DT, Mahon PB, Cai X, Kovacsics CE, Blackwell RA, Arad M, et al. Mood Disorder Susceptibility Gene CACNA1C Modifies Mood-Related Behaviors in Mice and Interacts with Sex to Influence Behavior in Mice and Diagnosis in Humans. Biological Psychiatry. 2010;68:801–810.

52. Dedic N, Pöhlmann ML, Richter JS, Mehta D, Czamara D, Metzger MW, et al. Cross-disorder risk gene CACNA1C differentially modulates susceptibility to psychiatric disorders during development and adulthood. Mol Psychiatry. 2018;23:533–543.

53. Lee AS, Ra S, Rajadhyaksha AM, Britt JK, De Jesus-Cortes H, Gonzales KL, et al. Forebrain elimination of cacna1c mediates anxiety-like behavior in mice. Mol Psychiatry. 2012;17:1054–1055.

54. Buske C, Gerlai R. Shoaling develops with age in Zebrafish (Danio rerio). Progress in Neuro-Psychopharmacology and Biological Psychiatry. 2011;35:1409–1415.

55. Geng Y, Peterson RT. The zebrafish subcortical social brain as a model for studying social behavior disorders. Dis Model Mech. 2019;12:dmm039446.

56. Buske C, Gerlai R. Early embryonic ethanol exposure impairs shoaling and the dopaminergic and serotoninergic systems in adult zebrafish. Neurotoxicology and Teratology. 2011;33:698–707.

57. Maaswinkel H, Zhu L, Weng W. Assessing Social Engagement in Heterogeneous Groups of Zebrafish: A New Paradigm for Autism-Like Behavioral Responses. PLOS ONE. 2013;8:e75955.

58. Lima-Ojeda JM, Mallien AS, Brandwein C, Lang UE, Hefter D, Inta D. Altered prepulse inhibition of the acoustic startle response in BDNF-deficient mice in a model of early postnatal hypoxia: implications for schizophrenia. Eur Arch Psychiatry Clin Neurosci. 2019;269:439–447.

59. Takahashi H, Hashimoto R, Iwase M, Ishii R, Kamio Y, Takeda M. Prepulse Inhibition of Startle Response: Recent Advances in Human Studies of Psychiatric Disease. Clin Psychopharmacol Neurosci. 2011;9:102–110.

60. Gottesman II, Gould TD. The Endophenotype Concept in Psychiatry: Etymology and Strategic Intentions. AJP. 2003;160:636–645.

61. Gould TD, Gottesman II. Psychiatric endophenotypes and the development of valid animal models. Genes, Brain and Behavior. 2006;5:113–119.

62. Kohl S, Heekeren K, Klosterkötter J, Kuhn J. Prepulse inhibition in psychiatric disorders – Apart from schizophrenia. Journal of Psychiatric Research. 2013;47:445–452.

63. Swerdlow NR, Light GA. Sensorimotor gating deficits in schizophrenia: Advancing our understanding of the phenotype, its neural circuitry and genetic substrates. Schizophrenia Research. 2018;198:1–5.

64. do Carmo Silva RX, Lima-Maximino MG, Maximino C. The aversive brain system of teleosts: Implications for neuroscience and biological psychiatry. Neuroscience & Biobehavioral Reviews. 2018;95:123–135.

65. Lee AS, De Jesús-Cortés H, Kabir ZD, Knobbe W, Orr M, Burgdorf C, et al. The Neuropsychiatric Disease-Associated Gene cacna1c Mediates Survival of Young Hippocampal Neurons. ENeuro. 2016;3.

66. Sykes L, Haddon J, Lancaster TM, Sykes A, Azzouni K, Ihssen N, et al. Genetic Variation in the Psychiatric Risk Gene CACNA1C Modulates Reversal Learning Across Species. Schizophrenia Bulletin. 2019;45:1024–1032.

67. Lisman J. Low-Frequency Brain Oscillations in Schizophrenia. JAMA Psychiatry. 2016;73:298.

68. Robertson MM, Furlong S, Voytek B, Donoghue T, Boettiger CA, Sheridan MA. EEG power spectral slope differs by ADHD status and stimulant medication exposure in early childhood. Journal of Neurophysiology. 2019;122:2427–2437.

69. Tiraboschi E, Martina S, Ent W van der, Grzyb K, Gawel K, Cordero-Maldonado ML, et al. New insights into the early mechanisms of epileptogenesis in a zebrafish model of Dravet syndrome. Epilepsia. 2020;61:549–560.

70. Bozarth X, Dines JN, Cong Q, Mirzaa GM, Foss K, Lawrence Merritt J, et al. Expanding clinical phenotype in *CACNA1C* related disorders: From neonatal onset severe epileptic encephalopathy to late-onset epilepsy. Am J Med Genet. 2018;176:2733–2739.

71. Uhlhaas PJ, Singer W. Abnormal neural oscillations and synchrony in schizophrenia. Nat Rev Neurosci. 2010;11:100–113.

72. Mäki-Marttunen T, Krull F, Bettella F, Hagen E, Næss S, Ness TV, et al. Alterations in Schizophrenia-Associated Genes Can Lead to Increased Power in Delta Oscillations. Cerebral Cortex. 2019;29:875–891.

73. Mäki-Marttunen T, Halnes G, Devor A, Witoelar A, Bettella F, Djurovic S, et al. Functional Effects of Schizophrenia-Linked Genetic Variants on Intrinsic Single-Neuron Excitability: A Modeling Study. Biological Psychiatry: Cognitive Neuroscience and Neuroimaging. 2016;1:49–59.

74. Mäki-Marttunen T, Lines GT, Edwards AG, Tveito A, Dale AM, Einevoll GT, et al. Pleiotropic effects of schizophrenia-associated genetic variants in neuron firing and cardiac pacemaking revealed by computational modeling. Transl Psychiatry. 2017;7:5.

75. Kumar D, Dedic N, Flachskamm C, Voulé S, Deussing JM, Kimura M. *Cacna1c* (Cav1.2) Modulates Electroencephalographic Rhythm and Rapid Eye Movement Sleep Recovery. Sleep. 2015;38:1371–1380.

76. Kantojärvi K, Liuhanen J, Saarenpää-Heikkilä O, Satomaa A-L, Kylliäinen A, Pölkki P, et al. Variants in calcium voltage-gated channel subunit Alpha1 C-gene (CACNA1C) are associated with sleep latency in infants. PLoS ONE. 2017;12:e0180652.

77. Saito R, Koebis M, Nagai T, Shimizu K, Liao J, Wulaer B, et al. Comprehensive analysis of a novel mouse model of the 22q11.2 deletion syndrome: a model with the most common 3.0-Mb deletion at the human 22q11.2 locus. Transl Psychiatry. 2020;10:35.

78. Namkung Y, Skrypnyk N, Jeong M-J, Lee T, Lee M-S, Kim H-L, et al. Requirement for the L-type Ca2+ channel α1D subunit in postnatal pancreatic β cell generation. J Clin Invest. 2001;108:1015–1022.

79. Platzer J, Engel J, Schrott-Fischer A, Stephan K, Bova S, Chen H, et al. Congenital deafness and sinoatrial node dysfunction in mice lacking class D L-type Ca2+ channels. Cell. 2000;102:89–97.

80. Langwieser N, Christel CJ, Kleppisch T, Hofmann F, Wotjak CT, Moosmang S. Homeostatic Switch in Hebbian Plasticity and Fear Learning after Sustained Loss of Cav1.2 Calcium Channels. Journal of Neuroscience. 2010;30:8367–8375.

81. Moosmang S, Haider N, Klugbauer N, Adelsberger H, Langwieser N, Müller J, et al. Role of Hippocampal Cav1.2 Ca2+ Channels in NMDA Receptor-Independent Synaptic Plasticity and Spatial Memory. Journal of Neuroscience. 2005;25:9883–9892.

82. Hofer NT, Tuluc P, Ortner NJ, Nikonishyna YV, Fernándes-Quintero ML, Liedl KR, et al. Biophysical classification of a CACNA1D de novo mutation as a high-risk mutation for a severe neurodevelopmental disorder. Mol Autism. 2020;11.

83. Pinggera A, Lieb A, Benedetti B, Lampert M, Monteleone S, Liedl KR, et al. CACNA1D De Novo Mutations in Autism Spectrum Disorders Activate Cav1.3 L-Type Calcium Channels. Biological Psychiatry. 2015;77:816–822.

84. Pinggera A, Mackenroth L, Rump A, Schallner J, Beleggia F, Wollnik B, et al. New gain-of-function mutation shows CACNA1D as recurrently mutated gene in autism spectrum disorders and epilepsy. Hum Mol Genet. 2017;26:2923–2932.

