## Supplementary materials for "Functional characterisation of single nucleotide variants of the psychiatric risk gene *cacna1c* in the zebrafish"

Banono et al.

### Supplementary Information

Supplementary Table 1

Supplementary Figures S1-6

### Supplementary Table

Table S1. The linearity data and calibration curve equations for the evaluated neuromodulators

| **Analyte** | Concentration Range (ng/10 µL injection) | Regression equation | Coefficient of determination ($R^2$) |
|---|---|---|---|
| Dopamine | 0.01-10 | y=2455983719x+6616427 | 0.9914 |
| GABA | 0.1-10 | y=1655578281x+5156982 | 0.9966 |
| Serotonin | 0.01-10 | y=1025282789x-6257299 | 0.9911 |
| Glutamate | 1-100 | y=12902625x-179259 | 0.9944 |

### Supplementary Figures

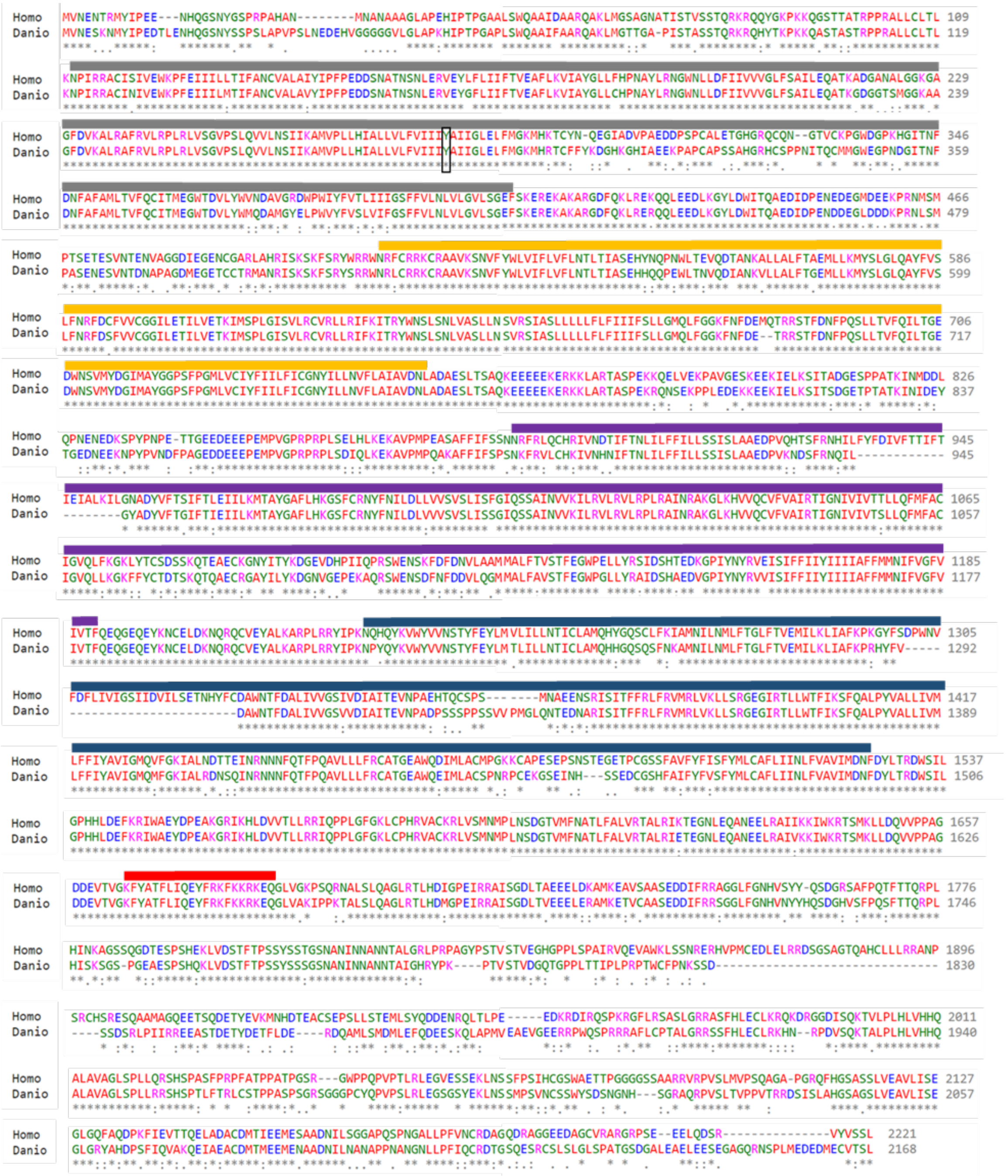

Figure S1. Alignment of CACNA1C protein sequence of humans (Homo) and zebrafish (Danio). Colour code of rectangular bars on top of aligning sequences: black – transmembrane domain I, yellow – transmembrane domain II, purple – transmembrane domain III, blue – transmembrane domain IV and red – IQ-domain. The black rectangular box represents the site of the *sa10930* mutation (Tyr292stop).

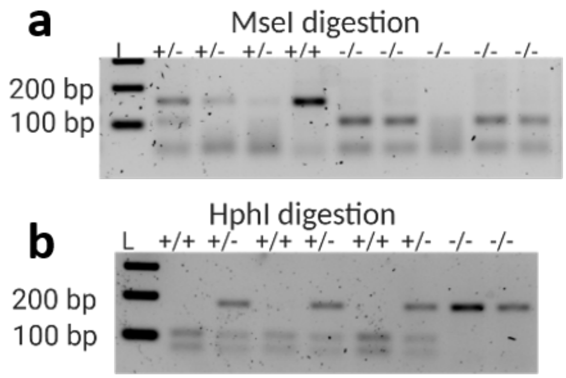


Figure S2. Genotyping of cacna1c mutants. a) Example gel electrophoresis of MseI digest of PCR products to determine the genotype of fish/larvae. MseI does not cut homozygous *sa10980* but cuts WT into two bands (70 and 119 bp), heterozygous *sa10930* into three bands (70, 119 and 196 base pairs). b) Example gel electrophoresis of HphI digest of PCR products to determine the genotype of fish/larvae. HphI does not cut homozygous *sa15296* but cuts WT into two bands (70 and 119 bp), heterozygous *sa15296* into three bands (70, 119 and 196 base pairs). Note the multiple bands in both MseI and HphI digested versus the undigested corresponding. -/-: WT siblings, +/-: heterozygous mutants, +/+: homozygous mutants.

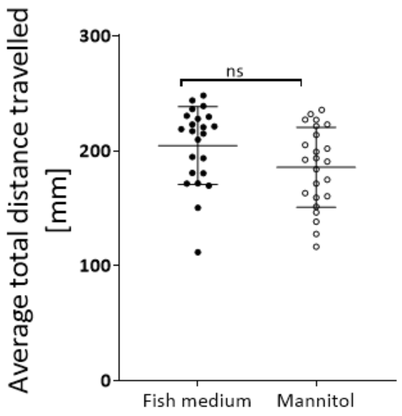


Figure S3. Mannitol does not affect the locomotor activity of wild type zebrafish larvae in comparison with fish medium. Data was analysed using unpaired student t-test and represented as Mean ± S.D. For the analysis, n = 23 and 24 in the fish medium and mannitol groups respectively.

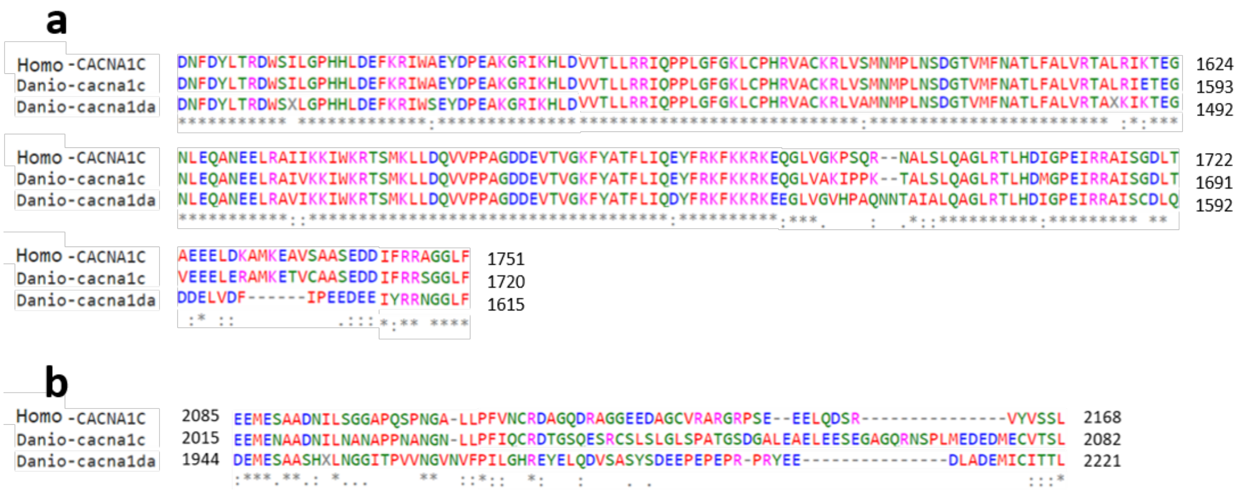


Figure S4. Alignment of sequences of antibodies used for western blotting and representative in situ staining. a) Anti-Ca$_V$1.2 fragment (MA5-277717). Possibility for unspecificity since the antigenic region is highly similar to zebrafish Cacna1da. b) Anti-Ca$_V$1.3 fragment (ab191038). Less possibility for unspecificity since the similarity of the antigenic region to zebrafish Cacna1c is very poor.

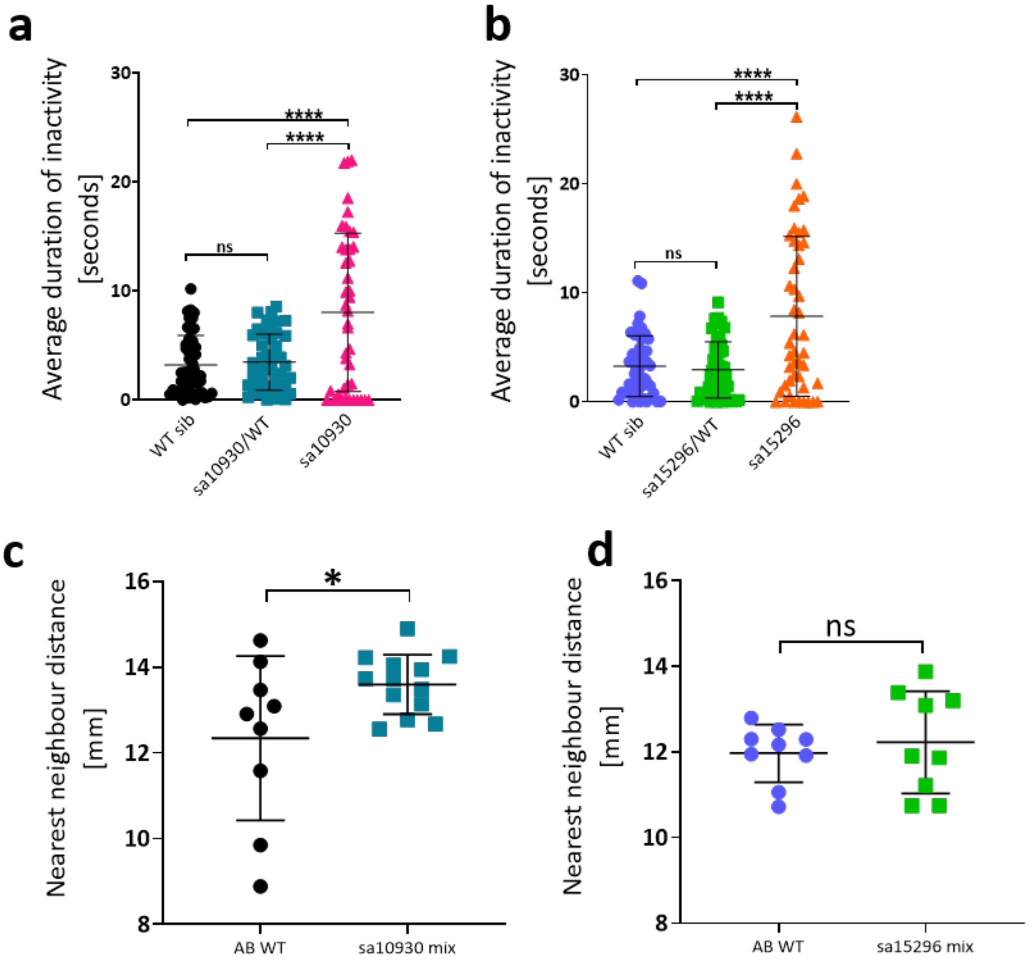


Figure S5. Behaviour. a-b) Duration of inactivity of *cacna1c* mutants versus their wild type siblings shown over 10 min in the locomotor activity test (a) *sa10930* mutant line and (b) *sa15296* mutant line. Data was analysed using Kruskal-Wallis one-way ANOVA test followed by Dunn's multiple comparison test and represented as mean ± S.D. For the analysis, n = 42-48, $^{**}p < 0.01$, $^{****}p < 0.0001$. c-d) Nearest-individual distance exhibited by a heterogeneous population of heterozygous mutants and their wild type siblings versus homogenous wild-type population. (c) *sa10930* mix and (d) *sa15296* mix. Data was analysed using unpaired Student *t*-test and represented as mean ± S.D. For the analysis, n = 9-13 (5 larvae/shoal), $^{*}p < 0.05$, $^{**}p < 0.01$.

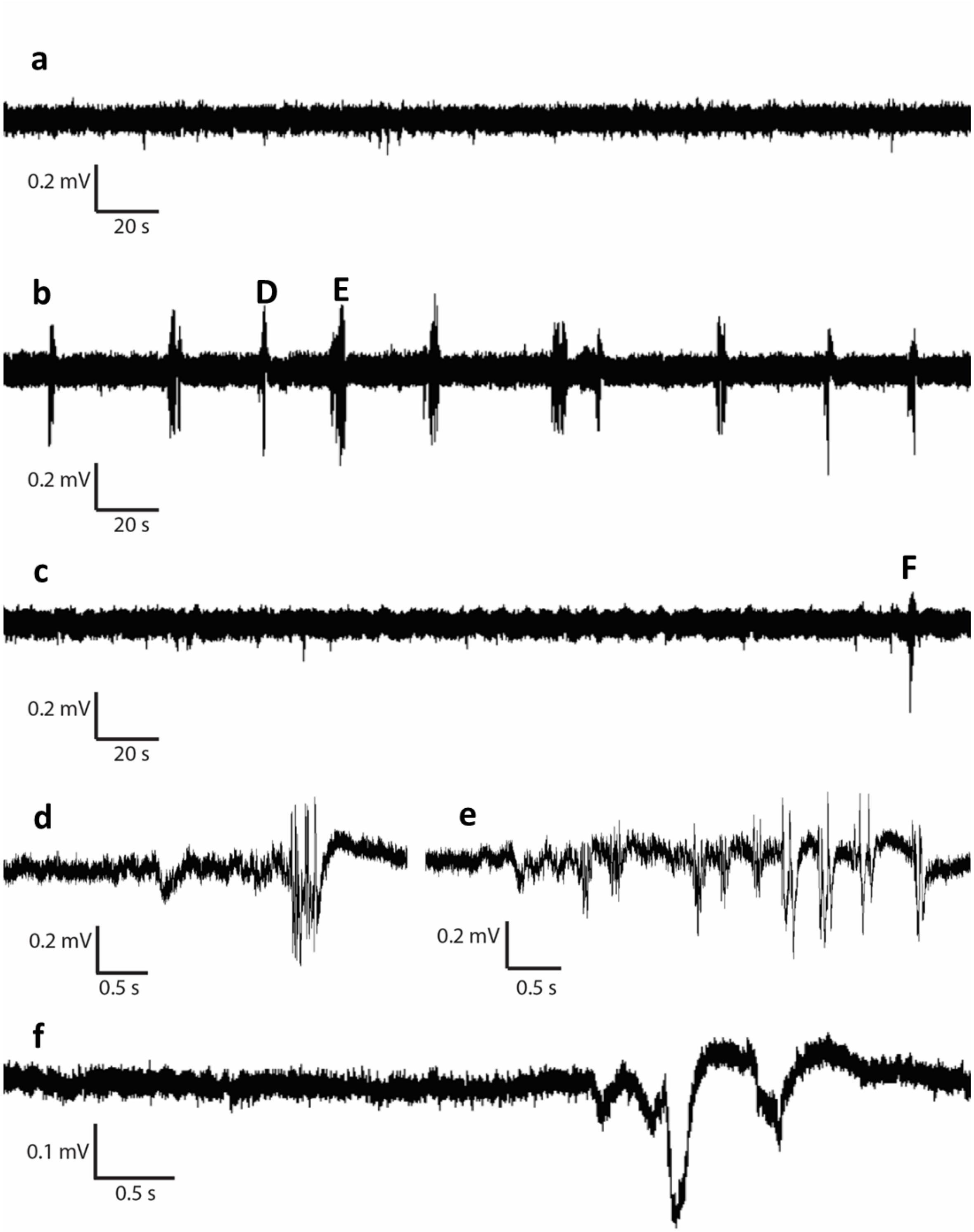


Figure S6. Representative LFP traces recorded from the optic tectum of 7 dpf zebrafish. Fragment of recording from a) AB WT, b) *sa15296* and c) *sa10930* larvae. d-f) are taken from the block letter locations in panels b and c.

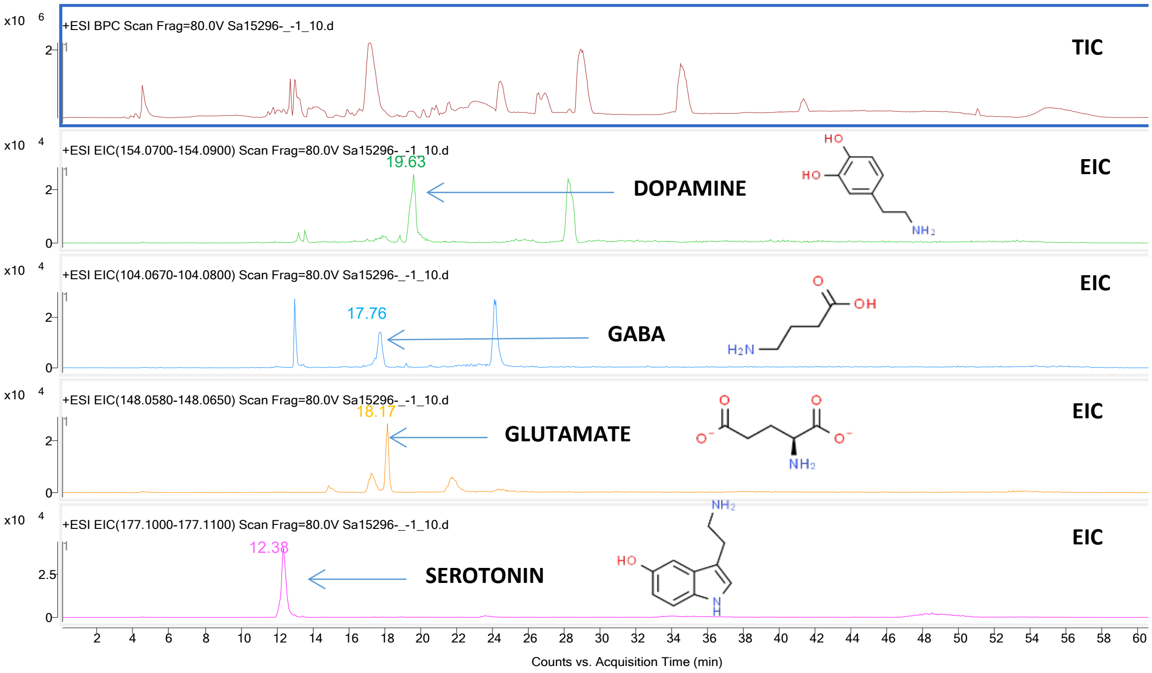


Figure S7. The mass chromatograms obtained from a sample extract, recorded in the positive ionization mode (TIC – the total ion chromatogram of the tested sample, EIC – the extracted ion chromatogram)
